# A natural broad-spectrum inhibitor of enveloped virus entry, effective against SARS-CoV-2 and Influenza A Virus in preclinical animal models

**DOI:** 10.1101/2022.02.16.480801

**Authors:** Rohan Narayan, Mansi Sharma, Rajesh Yadav, Abhijith Biji, Oyahida Khatun, Raju Rajmani, Pallavi Raj Sharma, Sharumathi Jeyasankar, Priya Rani, C. Durga Rao, Vijaya Satchidanandanam, Saumitra Das, Rachit Agarwal, Shashank Tripathi

**Affiliations:** Emerging Viral Pathogens Laboratory, Infosys Wing, Centre for Infectious Disease Research, Indian Institute of Science; Bengaluru, 560012, India; Department of Microbiology and Cell Biology, Division of Biological Sciences, Indian Institute of Science; Bengaluru, 560012, India; Centre for BioSystems Science and Engineering, Indian Institute of Science; Bengaluru, 560012, India; Molecular Biophysics Unit, Indian Institute of Science; Bengaluru, 560012, India; Department of Biological Sciences, School of Engineering and Sciences, SRM University, Andhra Pradesh, 522240, India

**Keywords:** Picolinic Acid, Broad-spectrum Antiviral, SARS-CoV-2, Influenza A virus, Preclinical Animal Model, Viral Entry, Membrane Fusion

## Abstract

The COVID-19 pandemic has highlighted the need for novel antivirals for pandemic management and preparedness. Targeting host processes that are co-opted by viruses is an attractive strategy for developing antivirals with a high resistance barrier. Picolinic acid (PA) is a byproduct of tryptophan metabolism, endogenously produced in humans and other mammals. Here we report broad-spectrum antiviral effects of PA against enveloped viruses, including Severe Acute Respiratory Syndrome Coronavirus-2 (SARS-CoV-2), Influenza A virus (IAV), Flaviviruses, Herpes Simplex Virus, and Human Parainfluenza Virus. We further demonstrate using animal models that PA is effective against SARS-CoV-2 and IAV, especially as an oral prophylactic. The mode of action studies revealed that PA inhibits viral entry of enveloped viruses, primarily by interfering with viral-cellular membrane fusion, inhibiting virus-mediated syncytia formation, and dysregulating cellular endocytosis. Overall, our data establish PA as a broad-spectrum antiviral agent, with promising preclinical efficacy against pandemic viruses SARS-CoV-2 and IAV.

## Introduction

Emerging and reemerging viral pathogens pose a great threat to public health due to their epidemic or pandemic potential. Novel strategies for developing broad-spectrum therapeutic interventions and repurposing in-pipeline drugs are crucial to tackling them (Meganck and Baric, 2021). SARS-CoV-2, the etiological agent of the currently ongoing Coronavirus Disease 2019 (COVID-19) pandemic (Zhu et al., 2020), has the world in its grip, and at the time of writing this manuscript, has caused over 410 million infections and > 5.8 million fatalities globally (https://coronavirus.jhu.edu/map.html). Influenza A viruses (IAV) on the other hand, cause yearly seasonal outbreaks, epidemics, and occasionally even global pandemics (Medina and García-Sastre, 2011). The dearth of effective antivirals for such viral diseases continues to take its toll both in terms of morbidity and mortality (Smed-Sörensen et al., 2018). SARS-CoV-2 is an enveloped virus with a single-stranded, positive-sense RNA genome, belonging to *Coronaviridae*, genus *betacoronavirus*. Its virion surface is decorated with surface Spike glycoprotein, which give the appearance of a crown (V’kovski et al., 2021). The SARS-CoV-2 spike binds to angiotensin-converting enzyme 2 (ACE2) receptors, and entry into host cells is facilitated by cleavage of spike by host proteases like transmembrane protease serine 2 (TMPRSS2) at the cell surface and cathepsin L in endosomes (Letko et al., 2020). Cleavage of spike results in S1 and S2 fragments; the former includes the N-terminal and receptor-binding domains (RBD), while the latter mediates virus-cell membrane fusion (Yu et al., 2022). Given the emergence of highly transmissible SARS-CoV-2 variants of concern and the ever-growing number of COVID-19 cases worldwide, there is an urgent need for effective antivirals to combat this disease.

Direct-acting antivirals target the viral components but generally are at a high risk of producing drug-resistant viral mutants, especially among RNA viruses (Sanjuán and Domingo-Calap, 2016). In contrast, cell-targeting drugs affect host factors that are obligate during the virus life cycle and ensuing pathogenesis (Lou et al., 2014). Targeting virus entry as an antiviral strategy has two main advantages - firstly, the virus lifecycle is inhibited early on, even before genomic replication can be initiated. Secondly, since virus entry mechanisms are limited, targeting one of these pathways can in principle inhibit a broad range of viruses that employ similar cellular pathways (Meganck and Baric, 2021, Mazzon and Marsh, 2019b). In the context of COVID-19, entry inhibitors could be used for pre-exposure prophylaxis and to limit the chances of SARS-CoV-2 antigenic drift (Chitsike and Duerksen-Hughes, 2021).

Picolinic acid (PA), also known as pyridine-2-carboxylic acid or 2-picolinic acid (PubChem, CID 1018), is a naturally occurring intermediate produced during the catabolism of Tryptophan via the Kynurenine pathway (Melillo et al., 1996, Grant et al., 2009). PA was recently shown to impact endosome biogenesis by directly inhibiting the E3 ubiquitin ligase UBR4 (Kim et al., 2018). Receptor-mediated endocytosis is a major pathway of viral entry into host cells, hence a target for broad-spectrum antiviral development (Mercer et al., 2010). Keeping this in mind, we examined the effect of PA on a range of human viral pathogens of clinical significance. In this study, we show that PA targets conserved mechanisms of viral entry to exert a broad-spectrum antiviral effect. Detailed investigation of the mechanism of action (MoA) revealed that PA act primarily by inhibiting viral and cellular membrane fusion, with potential additional effect due to interference with virus-mediated host cell syncytia formation and cellular endocytic processes. Unsurprisingly, PA was found to be effective against a wide range of enveloped viruses, with limited or no effect against non-enveloped viruses. More importantly, PA showed promising preclinical antiviral efficacy against SARS-CoV-2 in the Syrian hamster model and IAV in BALB/c murine model. The *in vivo* effects were observed in both prophylactic as well as therapeutic regimen and oral as well as intraperitoneal administration of PA at non-toxic levels. Overall, this study paves the way for clinical development and use of picolinic acid as a broad-spectrum antiviral, which can help in combating the COVID-19 pandemic as well as other emerging and re-emerging viruses.

## Results

### Picolinic exhibits *in vitro* antiviral activity against SARS-CoV-2, IAV, and a range of other enveloped viruses

Picolinic acid is an organic compound, produced naturally through the catabolism of Tryptophan (Fig 1A). It has been reported to interfere with the maturation of early endosomes (Kim et al., 2018). We reasoned that since the endocytic process is co-opted for cellular entry by a range of viruses, PA must interfere with their infection. To test this, we first examined the *in vitro* antiviral efficacy of PA against IAV. To this end, we tested the PA dose-dependent effect on A/Puerto Rico/8/1934 H1N1 IAV (PR8) replication in Madin-darby canine kidney (MDCK) cells and observed a dose-dependent decrease in infectious virus counts at 48hr post-infection (p.i), with maximum inhibition seen in the presence of 2 mM PA (IC50=1.83 mM) (Fig 1B). We further went on to test the antiviral activity of PA against the 2009 H1N1 ‘Swine Flu’ pandemic IAV (Cal/09) and observed consistent inhibition of viral replication (Fig 1C). Next, we began to explore the potential broad-spectrum antiviral activity of PA against other important human viral pathogens. To this end, we tested PA dose-dependent effect on Hong Kong/VM20001061/2020 (SARS-CoV-2 Hong Kong) replication in, human HEK293T-ACE2 cells with exogenous ACE2 receptor expression. We observed ∼2-log_10_ reduction in viral RNA (vRNA) load in the presence of 2 mM PA, which was non-toxic to cells, with an IC50 of 0.87 mM (Fig 1 D). This effect was further tested against SARS-CoV-2 variants of concern (VOC) including Alpha, Beta, Gamma, and Delta VOCs and the antiviral effect of PA was consistently observed (Fig 1E). These findings were validated in a similar experiment using primate origin Vero E6 cells, where 2 mM PA treatment reduced SARS-CoV-2 vRNA copy by ∼2-log_10_ with IC50 of 0.59 mM (Fig 1F). Also, the antiviral activity of PA was found to be conserved against SARS-CoV-2 VOCs in this cell line as well (Fig 1G). We also validated the anti-SARS-CoV-2 activity of PA in Calu-3 cells which are of human lung tissue origin and endogenously express ACE2 receptor, where a similar ∼2 log_10_ reduction in viral RNA copy numbers was observed in PA treated cells (Sup Fig 1A). Next, we further expanded the test of broad-spectrum antiviral activity of PA, now against Flaviruses, another clinically important group of human viruses. For this, we used human lung carcinoma A549 cells and established that PA at 2 mM concentration was non-toxic in them (CC50 - 9.17 mM) (Sup Fig 1B). Multi-cycle infection in A549 cells infected with DENV and Zika reporter viruses expressing luciferase showed >90% inhibition in the presence of 2mM PA (Fig 1 H, I). The anti-Flavivirus effects of PA were supported by inhibition of Japanese encephalitis Virus (JEV) clinical strain P20778, and Asian lineage Zika virus (ZIKV) Cambodia strain as shown by western blot analysis of Flavivirus envelope protein (Sup Fig 1 C-F). We also tested the antiviral activity of PA against the Human parainfluenza virus (HPIV-3) and Herpes simplex type 1 virus (HSV-1), expressing luciferase reporter, and observed 80-90% reduction at 2 mM PA concentration (Fig 1 J, K). With this set of experiments, the *in vitro* broad-spectrum antiviral activity of PA against a range of enveloped viruses was confirmed.

**Figure 1:**
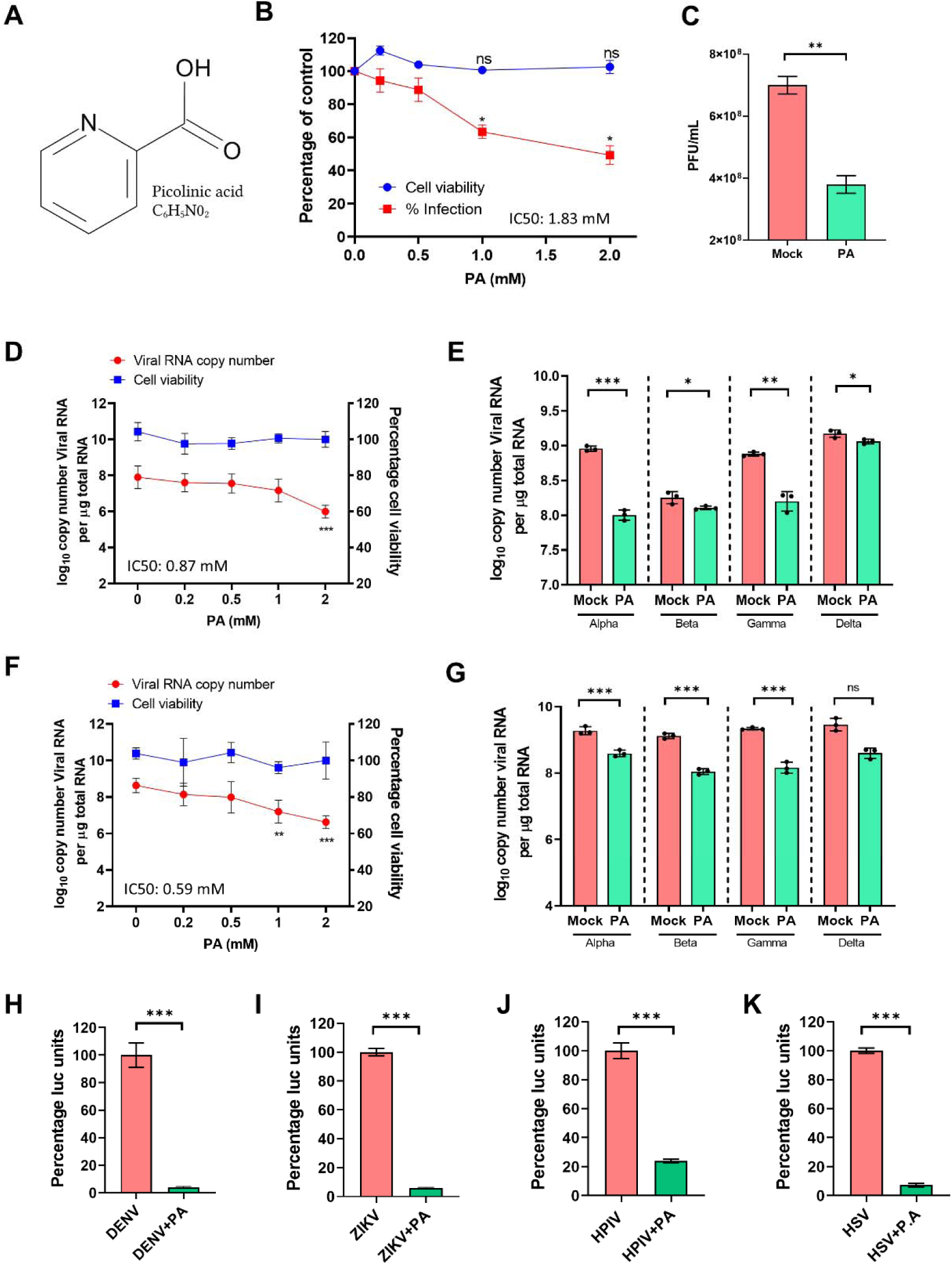
Picolinic Acid exhibits broad-spectrum antiviral activity. (A) Chemical structure of Picolinic acid (B) MDCK cells pre-treated with increasing concentrations of PA were infected with 0.01 MOI of PR8 IAV and 48hr p.i, virus from supernatants were quantified by plaque assay. Results were plotted along with cell viability measured from uninfected cells treated with PA. (C) MDCK cells pre-treated with 2 mM PA were infected with Cal/09 IAV and 48hr p.i, virus from supernatants were quantified by plaque assay. (D-E) HEK293T-ACE2 cells were pre-treated for 3hr with increasing doses of PA as indicated, infected with 0.1 MOI of either (D) SARS-CoV-2 Hong Kong or (E) four SARS-CoV-2 variants of concern. Cells were collected 48hr p.i, viral RNA copy estimated by qRT PCR, and corresponding cell viability of uninfected drug-treated cells were plotted. (F) and (G) show datasets corresponding to Vero E6 cells infected with 0.001 MOI for all viruses as mentioned above. (H-K) A549 cells pre-treated with 2 mM PA were infected with different luciferase reporter viruses as indicated. Cells were harvested 48hr p.i for luciferase assay and data normalized with untreated control are shown. *p < 0.05; **p < 0.01; ***p < 0.001; ns - not significant, using two-tailed unpaired t-test or one-way ANOVA with Dunnett’s multiple comparison test, wherever necessary. Error bars represent mean ± standard deviation.

### Picolinic acid treatment inhibits Influenza A virus replication and pathogenesis in a preclinical murine model

Given the broad-spectrum activity of PA across multiple viral pathogens and the urgent need for antiviral options against pandemic viruses, we went on to test *in vivo* efficacy of PA, where animal models of viral pathogenesis were available. To start, we first tested the preclinical efficacy of PA against IAV in the BALB/c murine model. We first conducted PA toxicity studies in BALB/c mice using 20 and 100 mg/kg body weight of drug administered through the intraperitoneal (IP) or oral route and found the drug non-toxic at 20 mg/kg dose in both IP and oral routes of administration (Fig 2A, B). At 100 mg/kg dose it was still well tolerated orally however through IP route was toxic, as indicated by ∼10% loss of weight over 9 days (Fig 2A, B). Based on this, 20 mg/kg PA was used for all treatment regimens via both IP and oral routes in all animal model studies. To test antiviral efficacy, we challenged BALB/c mice with intranasal inoculation of 50 PFU PR8 IAV and subjected them to either prophylactic or therapeutic treatment with PA, with oral or IP administration of the drug (Fig 2C). In all cases, PA treatment significantly improved animal survival, which was 100% for all treated groups except IP prophylactic group with 80% survival, against 20% survival in the untreated control group (Fig 2D). Consistent with this, PA treatment in all formats also prevented body weight loss significantly as compared to the untreated group, over 7 days (Fig 2E). The antiviral effects of PA against IAV were also reflected in reduced infectious virus count in mice lungs, with the better reduction seen in the prophylactic group (Fig 2F). Histopathological examination of the lung tissue from infected animals also showed reduced pathology features (vascular and alveolar infiltration of cells and interstitial pneumonia) in the case of drug-treated groups as compared to untreated control (Fig 2G, H). Overall, both IP and oral treatment with PA displayed promising *in vivo* antiviral effects against IAV, with better outcomes in prophylactic when compared to therapeutic regimen.

**Figure 2:**
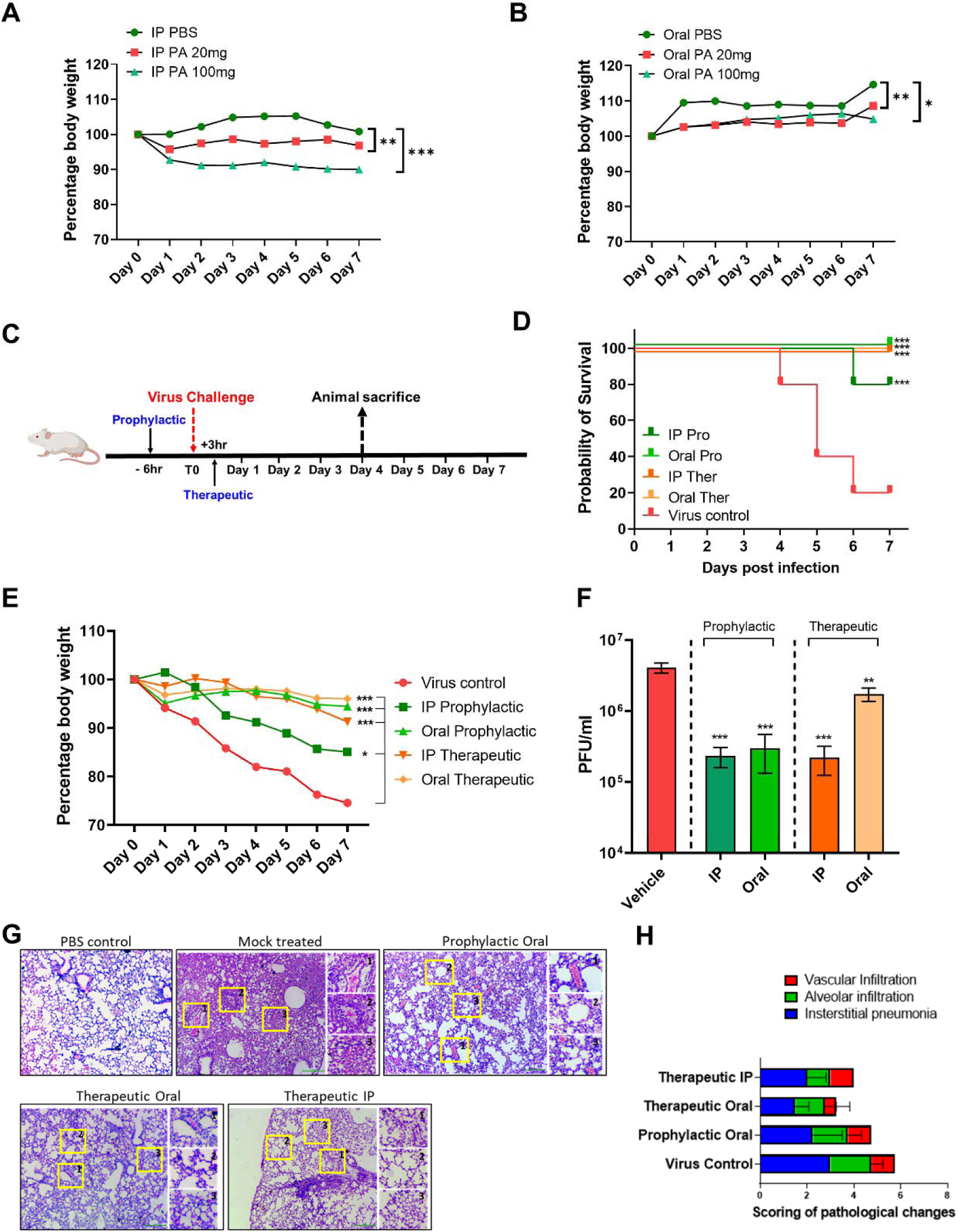
Picolinic Acid mitigates IAV replication and pathogenesis *in vivo*. (A-B) Toxicity results as shown by bodyweight changes over 9 days post-treatment with 20 or 100mg/kg PA delivered via (A) IP and (B) oral routes. Data are from 1 experiment and presented as the mean percentage of bodyweight at Day 0, n= 2-3 per group. One-way ANOVA Mixed-effects analysis with Dunnett’s multiple comparisons against IP or Oral PBS groups were performed.*p < 0.05; **p < 0.01; ***p < 0.001.. ns - not significant. (C) Schematic showing infection schedule. The prophylactic and therapeutic treatment used administration of PA 6hr prior and 3hr post-infection respectively, survival was monitored up to day 4 p.i and bodyweight loss for remaining animals till day 7. (D) Survival of mice (n=5 per group) was monitored over the course of 7 days p.i for control and treated groups. Survival curves from 1 experiment is shown, log-rank (Mantel–Cox) test, ***p < 0.001. (E) Body weight loss of animals monitored up to 7 days p.i. Data are from 1 experiment and presented as the mean percentage of bodyweight at Day 0, n=2-5 per group. p values were calculated by One-way ANOVA with Dunnett’susing multiple comparisons.*p < 0.05; ***p < 0.001. after mixed-effects analysis. *p < 0.05. (F) Plaque assay quantification of infectious virus titer from lungs. (G) Histology images for treatment groups including controls. p values were calculated from one-way ANOVA with Dunnett’s multiple comparison test, **p < 0.01; ***p < 0.001. Error bars represent mean ± standard deviation. (G) Histology images for treatment groups including controls. (H) Scoring of clinical pathology was done based on the following criteria, namely (1) vascular infiltration, (2) alveolar infiltration, and (3) interstitial pneumonia. Scoring was done based on the severity on a scale of 1-4 (1-mild, 2-moderate, 3-severe, 4-very severe). An overall score was given by accumulating the total scores for each criterion.

### Picolinic acid treatment inhibits SARS-CoV-2 replication and pathogenesis in a preclinical Syrian hamster model

With encouraging results of *in vivo* efficacy against IAV, we next tested the same against SARS-CoV-2. For this, we used the Syrian Golden hamster model and an optimized protocol for testing antivirals against SARS-CoV-2 developed by our group (Fig 3A) (Biji et al., 2021). We observed that IP administration of 20 mg/kg PA prophylactically and therapeutically, resulted in a reduction of lung vRNA load on D3 post-infection by ∼3 and ∼1 order of magnitude respectively (Fig 3B). This was accompanied by a significant reduction in lung inflammation and body weight loss in PA-treated groups (Fig 3C, D). Further, similar protective efficacy effects of PA were obtained upon oral administration, including the inhibitory effect on lung vRNA load, inflammation, and animal body weight (Fig 3E, F, G). Evidence from histopathology analysis of lung tissue sections clearly showed decreased pathology, supported by a marked reduction in alveolar edema and cellular infiltration of lungs in PA treated groups (Fig 3 H, I). Overall, results indicate PA exhibits antiviral activity against SARS-CoV-2 in the preclinical Syrian hamster model, with the best outcome in the case of prophylactic treatment.

**Figure 3:**
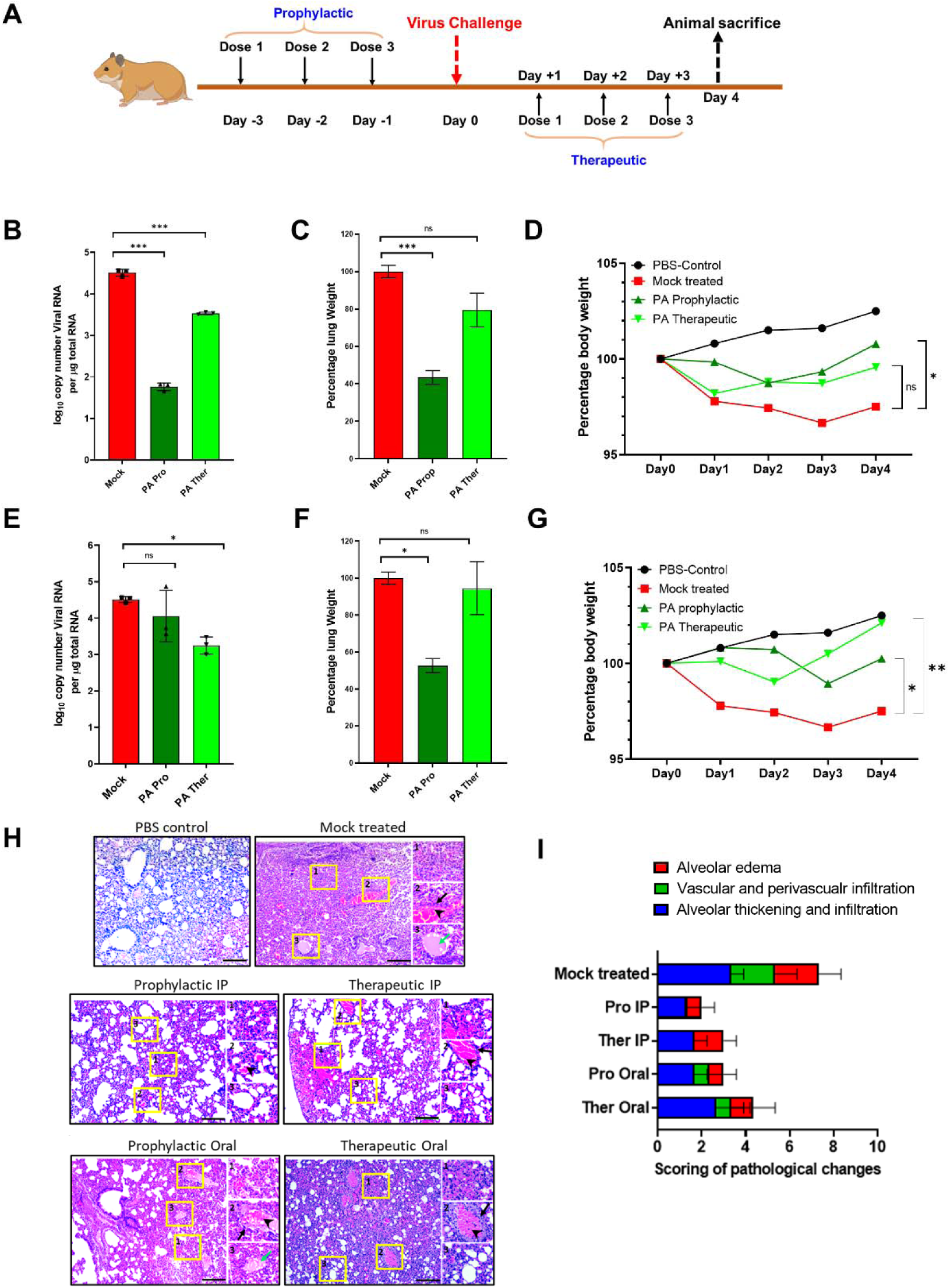
Picolinic Acid mitigates SARS-CoV-2 replication and pathogenesis *in vivo*. (A) Schematic for PA treatment in hamsters using prophylactic and therapeutic regimens is shown. Prophylactic treatment of hamsters involved administration of PA at 1,2 and 3 days prior to infection, followed by virus challenge at day 0. The therapeutic treatment used dosage during 1,2 and 3 dpi. In both cases, animals were sacrificed 4dpi. (B-D) Results for the administration of PA via IP route showing (B) lung vRNA copy number, total lung weight, and (D) bodyweight of animals up to 4 dpi. (E-G) Corresponding data for oral administration of PA showing (E) lung viral RNA copy number, (F) total lung weight, and (G) bodyweight of animals. qRT PCR and lung weight data shown are from 1 experiment (n=3 per group). Bodyweight data are from 1 experiment and presented as the mean percentage of bodyweight measured at Day 0, n= 2-5 per group. In all cases, One-way ANOVA with Dunnett’s multiple comparisons were performed.*p < 0.05; **p < 0.01. (H) Histology images for all treatment groups including mock-infected and healthy controls. (I) Clinical scoring was based on the following criteria and labelled within inset images as (1) alveolar edema, (2) vascular and perivascular infiltration and (3) alveolar thickening and infiltration. Black arrows indicate vascular infiltration, arrowheads show perivascular infiltration, green arrow show alveolar edema. Scoring was done based on the severity on a scale of 1-4 (1-mild, 2-moderate, 3-severe, 4-very severe). An overall score was given by accumulating the total scores for each criterion.

### Time of addition studies reveals inhibition of virus entry as the mechanism of antiviral action for picolinic acid

With broad-spectrum antiviral activity and *in vivo* effects against IAV and SARS-CoV-2 established, we next aimed to understand the mechanism of action (MoA) of PA. For this, we used SARS-CoV-2 and IAV as test viruses and conducted time of addition assays (ToA), wherein cells were treated with the drug at different time points before, during, and after infection to identify the steps of inhibition during a single cycle of virus infection (Aoki-Utsubo et al., 2018). In case of SARS-CoV-2 infection, pre-treatment of HEK293T-ACE2 cells with 2 mM PA for 3hr (−3hr) mitigated virus infection by >80% as shown by immunofluorescence assay (IFA) images and quantification of cells positive for virus spike protein, as well as quantification of spike protein by western blotting (Fig 4 A, B, C). Similar effects were also seen when the virus and drug were incubated for 1hr before infection. However, without pre-treatment of cells, when PA was added at the time of infection (T0), the only limited reduction was observed in Spike positive cells or protein levels (Fig 4 A, B, C). To test the effects of PA on post-entry stages of the virus replication cycle, the drug was added 6hr post-infection and collected 3hr later. Here, no inhibition of virus infection was observed, either on Spike positive cells or protein levels (Fig 4 D, E, F). A similar time of addition experiment was conducted on Vero E6 cells, where pre-treatment (−3hr) with PA inhibited virus infection (Fig 4 G, H, I). However, introduction of drug at the time of infection (T0) or post-infection (T+6) did not affect virus infection (Fig 4 J, K, L). These results confirmed that PA affects the early stages of SARS-CoV-2 infection in the host cells and doesn’t affect later events of the viral life cycle. To confirm the effect of PA on virus entry, we utilized SARS-CoV-2 spike pseudotyped particles expressing luciferase reporter, which recapitulate exclusively the virus entry step (Crawford et al., 2020). Here, in the presence of increasing PA concentrations, we observed a dose-dependent effect, with 90% inhibition of Luciferase expression at 2 mM concentration in Vero E6 cells (Fig 4 M). A similar effect of PA was observed in Spike pseudotyped particle entry assay on Calu-3 cells (Fig 4 N). These results confirmed that the antiviral action of PA against SARS-CoV-2 involves inhibition of viral entry in the host cell. It has been reported that SARS-CoV-2 induces cell-cell fusion and syncytia formation, primarily through the fusogenic activity of Spike (Yu et al., 2022). These events involve similar Spike activity that facilitates membrane fusion events during viral entry (Tang et al., 2020). Hence, we tested whether PA has inhibitory activity against syncytium formation by SARS-CoV-2 Spike protein. To this end, Vero E6 cells transfected with a plasmid expressing SARS-CoV-2 spike protein were treated with increasing concentrations of PA, and 24hr later, fixed and labeled with wheat germ agglutinin (WGA) to mark the cell membrane and DAPI to mark the nucleus. Results showed a dose-dependent inhibition of virus spike-induced syncytia formation. Treatment with 2 mM PA resulted in 80% inhibition of syncytia, as shown by IFA images and quantification of syncytia (Fig 4 O, P). Taken together these data confirm that the primary action of PA against SARS-CoV-2 is inhibition of viral entry, however in a multicycle experiment or in vivo it may have an additional effect by inhibiting syncytia formation.

**Figure 4:**
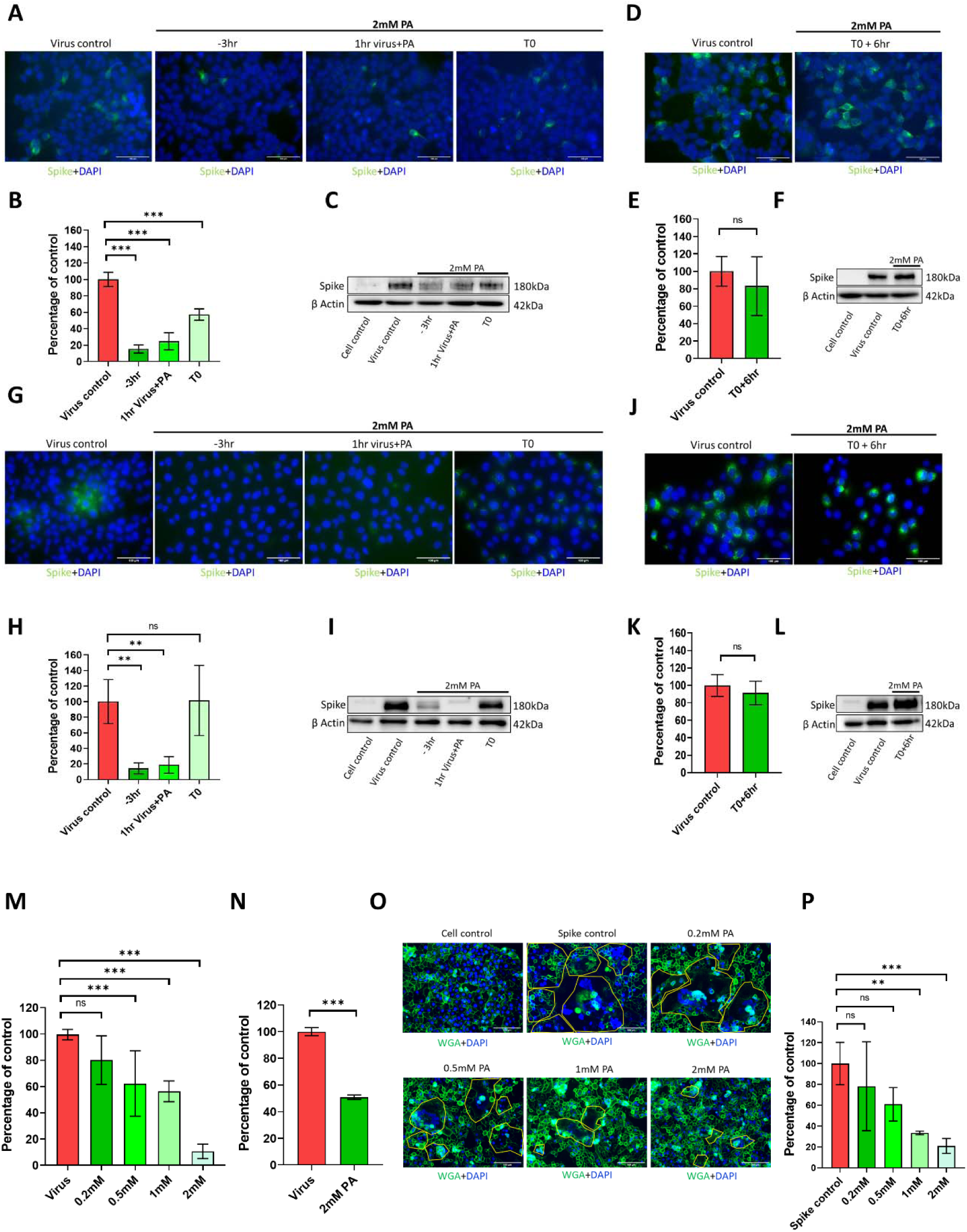
Picolinic Acid inhibits SARS-CoV-2 entry and Spike mediated Syncytia formation. (A-C) HEK293T-ACE2 cells were either pre-treated for 3hr with 2mM PA, infected with 10 MOI SARS-CoV-2 Hong Kong in presence of the drug and collected 3hr later (−3hr), or virus and drug were incubated together for 1hr and used for infection (1hr virus+PA), or cells were treated at the time of infection (T0). Microscopy images of infected cells are shown in (A), quantification of infected cells labeled using anti-spike antibody is shown in (B), and corresponding western blot analysis of above conditions in (C). (D-F) HEK293T-ACE2 cells were first infected with 10 MOI SARS-CoV-2 Hong Kong, PA treatment was done 6hr p.i (T0+6hr) and cells were collected a further 3hr later. Microscopy images with quantification and western blot analysis are shown in (D), (E), and (F). (G-I) and (J-L) show similar datasets for Vero E6 cells infected with 10 MOI SARS-CoV-2 Hong Kong. (M) HEK293T-ACE2 cells were pre-treated with increasing doses of PA as indicated, infected with SARS-CoV-2 spike pseudotyped particles in the presence of the drug and harvested 60hr later. Results show firefly luciferase values normalized to virus control. (N) Similar experimental conditions were followed for Calu-3 cells but using only 2mM PA treatment. Results are shown in (N). (O) Vero E6 cells were transfected with a plasmid expressing SARS-CoV-2 spike and 3hr later treated with increasing concentrations of PA as indicated. 24hr later, cells were fixed and labeled with WGA and DAPI to label cell membrane and nuclei respectively. The area of syncytia was quantified using ImageJ/Fiji and plotted as a percentage of untreated control (P). **p < 0.01; ***p < 0.001; ns - not significant, using two-tailed unpaired t-test or one-way ANOVA with Dunnett’s multiple comparison test, wherever necessary. Error bars represent mean ± standard deviation.

### Picolinic acid inhibits virus entry by disrupting the viral membrane and interfering with viral-cellular membrane fusion

To further explore, whether PA act by similar MoA against other viruses, we performed ToA experiments using IAV. Here also pretreatment (−3hr) of A549 cells with PA reduced PR8 IAV infection by 80%, as measured by reduction in IAV nucleoprotein (NP) positive cells at 3hr post-infection, as well as NP protein level in western blot (Fig 5 A, B, C). Also, similar to SARS-CoV-2, IAV incubation with drug for 1hr and subsequent infection, showed 50% inhibition of entry, as observed by IFA and western blotting for NP (Fig 5 A, B, C). No differences were observed when PA was added 6hr post-infection (Fig 5 D, E, F). In the case of SARS-CoV-2, PA inhibited the fusogenic activity of Spike protein. For IAV fusion of viral and cellular membrane takes place inside endosomal compartment due to pH-dependent conformational changes triggered in the hemagglutinin (HA) protein (Matlin et al., 1981) To test the effect of PA on IAV viral-cellular membrane fusion we used an established protocol (Hoffmann et al., 2018) where virions are labeled with a lipophilic fluorescent dye and are allowed to attach to target cells at low temperature, followed by raising the temperature to 37 °C to trigger viral endocytosis and entry. In intact virion, the dye signal is quenched, however upon viral entry, the dye is de-quenched, and the fluorescence signal is produced, which indicates fusion of viral and endocytic membrane (Fig 5G). In a similar assay, we used R18 dye-labeled PR8 IAV virions to infect PA-treated MDCK cells on ice. Subsequently, the temperature was raised to 37°C, and the increasing fluorescence signal associated with virus-endosome fusion during virus entry was quantified over a period of 90 minutes. Results showed that cells pretreated with 2mM PA were all able to inhibit virus-entry-associated viral-endocytic membrane fusion and increase in fluorescence caused by R18 dye dispersal. This activity of PA was similar to well-known pH-dependent endocytic viral entry inhibitors ammonium chloride (NH_4_Cl) and Chloroquine diphosphate (CQ) (Yoshimura and Ohnishi, 1984) To confirm whether, in the case of IAV, PA treatment affects late events of the replication cycle, a mini replicon assay where co-transfection of IAV polymerase components drives expression of the Luciferase reporter gene was performed. Results showed that 2mM PA treatment had no impact on IAV polymerase activity at 8 and 12hr post-treatment (Fig 5I). These data established that PA acts against IAV by inhibiting viral entry, specifically the fusion of viral and endosomal membranes, and has no impact on late events of IAV replication or polymerase activity. To further understand the mechanistic basis of viral-cellular membrane fusion inhibition, we used transmission electron microscopy (TEM) imaging of PA-treated PR8 IAV virions to examine potential effects on viral structural integrity. Results clearly showed severe disruption of viral envelope compared to untreated control. These results indicate that PA treatment inhibits viral and cellular membrane fusion, primarily by affecting the integrity of the viral membrane, which is likely to be common MoA of PA against enveloped viruses.

**Figure 5:**
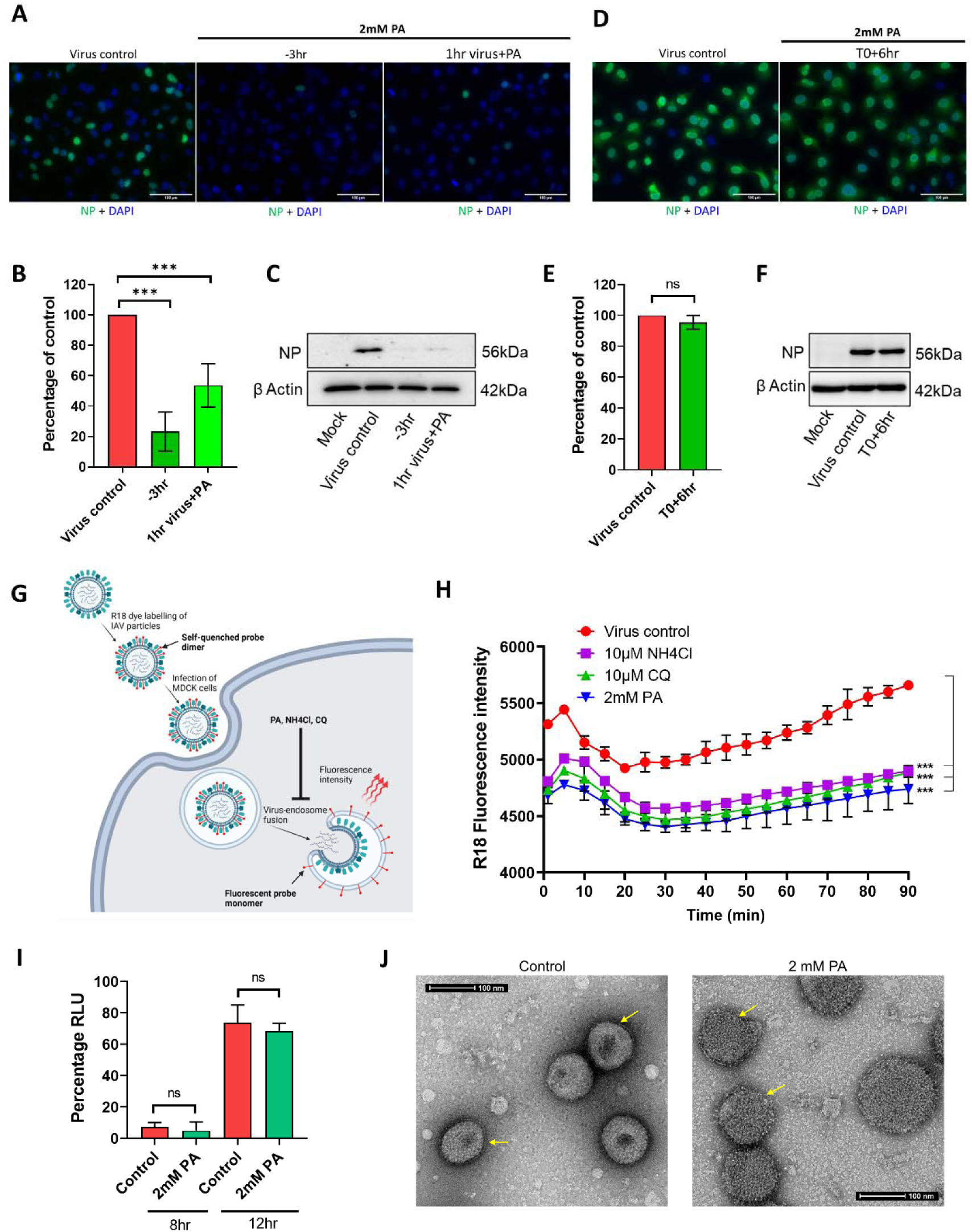
Picolinic Acid inhibits IAV entry by interfering with Viral-Cellular endocytic membrane fusion and affecting viral membrane integrity. (A-C) A549 cells were either pre-treated for 3hr with 2mM PA, infected with 2 MOI PR8 WT in presence of the drug, and collected 3hr later (−3hr) or the virus was incubated with 2mM PA for 1hr and used for infection (1hr virus + PA). Microscopy data for A549 cells labeled with influenza virus nucleocapsid protein are shown in (A), quantification of NP positive cells is shown in (B), and (C) shows corresponding western blot data. (D-F) A549 cells were first infected with 2 MOI PR8 WT, PA treatment was done 6hr p.i (T0+6hr) and cells were collected a further 3hr later. Microscopy images with quantification and western blot analysis are shown in (D), (E), and (F). ***p < 0.001; ns - not significant using two-tailed unpaired t-test or one-way ANOVA with Dunnett’s multiple comparison test where applicable. Error bars represent mean ± standard deviation. (G) Pictorial representation of virus-endosome membrane fusion assay using R-18 labeled IAV particles. After infection of MDCK cells, the labeled virus particles enter endosomes, and upon pH-dependent fusion of virus and endosomal membranes, results in the de-quenching of R-18 probe dimers and subsequent increase in fluorescence intensity. Created with Biorender. (H) A549 cells were pre-treated with either 2mM PA, 10µM NH4Cl, or 10µM CQ, infected with R18 labeled PR8 WT virus on ice, transferred to a plate reader at 37°C and fluorescence intensity measurements were acquired at 10 min intervals. ***p < 0.001, using one-way ANOVA with Dunnett’s multiple comparison at 90 min time point. (I) HEK293T cells were transfected with plasmids expressing influenza virus PA, PB1, PB2, NP, and NP-firefly luc, along with pRLTK. 2mM PA was added 3hr post-transfection and cells were harvested 8 and 12hr later. Results show luciferase units normalized to untreated control. ns - not significant using two-tailed unpaired t-test. Error bars represent mean ± standard deviation. (J) Concentrated PR8 WT virus particles were incubated with vehicle control or 2mM PA for 3hr, mounted on copper grids, and processed for TEM imaging. Arrows indicate differences in the integrity of viral double-layered membranes in control and treated conditions.

### Picolinic acid exhibits minimal to no antiviral activity against non-enveloped viruses, including bacteriophages

So far, we had established broad-spectrum antiviral activity of PA against a battery of enveloped viruses and mapped its MoA to inhibition of viral entry, primarily by interfering with viral and cellular membranes. However, the question remained that if antiviral effects of PA were analogous to disruption of virus-cell membrane fusion during entry, will the drug be equally effective against non-enveloped viruses as well? To this end, we tested a range of non-enveloped viruses starting with Coxsackie virus B3. Entry experiments performed in HeLa cells showed that pre-treatment of cells, incubation of virus with the drug before infection, and treatment at the time of infection, did not have any effects on virus entry, as seen by western blot data for virus VP1 protein (Fig 6 A). Even in the multicycle infection experiment, PA was ineffective against the Coxsackie virus, as reflected in plaque assay-based measurement of infectious virus count in cell supernatant (Fig 6 B). Thereafter, we examined the effect of PA on Rhesus monkey rotavirus (RRV) infection in HEK293T cells. Here also pre-treatment of cells with PA did not result in any differences in virus infection, as shown by IFA images and quantification of VP6 positive cells (Fig 6 C, D). We then proceeded by evaluating PA effects on non-enveloped Adeno associated virus particles expressing eGFP (AAV6-eGFP). HEK293T cells infected with increasing volumes of the AAV6-eGFP preparations in the presence of 2mM PA resulted in a 5-10% decrease in GFP positive cells in all conditions tested (Fig 6 E). In a similar experimental setup but using non-enveloped Adeno virus 5 particles expressing eGFP (Ad5-eGFP) for 24hr infection, no differences were observed in GFP positive HEK293 cells count, between PA treated and untreated conditions (Fig 6 F). All the viruses tested so far infect humans or higher vertebrates. We examined further whether PA has antiviral activity against bacteriophages. For this, we used TM4 mycobacteriophages, which are non-enveloped, and tested the effect of PA on their infection in *Mycobacterium smegmatis* bacteria. To rule out any toxicity of PA treatment on *M. smegmatis*, the bacteria were grown in 7H9 broth in presence of a range of PA concentrations. Optical Density measurement of the culture at 600 nm for 24hr showed 1mM PA to be the highest non-toxic concentration (Fig 6 G). Next, to test for the antiviral activity, 1mM PA was either added at the start of the experiment or 3hr prior to infection of *M. smegmatis* with TM4 phage. In either case, there was no protection from TM4 mycobacteriophage infection-induced bacterial cell death (Fig 6H). This indicated that PA does not affect the entry of non-enveloped bacteriophages in bacteria. Although non-enveloped viruses tested in our studies do not engage in viral-cellular membrane fusion, they do depend on cellular endocytosis for entry into host cells (Meier and Greber, 2004, Marjomäki et al., 2015, Bartlett et al., 2000). PA has been reported previously to interfere with endocytic maturation in mammalian cells (Kim et al., 2018). To confirm this, we looked at the effect of PA treatment on transferrin uptake and recycling, which is an established measure of the endocytic process (Widera et al., 2003, Lakadamyali et al., 2006) Specifically, we used fluorescent transferrin conjugates (Tf 647) to pulse A549 cells for 1hr, and measured Tf647 uptake and recycling by IFA and FACS. We observed that the presence of 2mM PA caused mislocalization of Tf647 loaded endocytic vesicles, as they were more dispersed throughout the cytoplasm as compared to perinuclear in the case of control cells (Sup Fig 2A, B). However, PA treatment did not affect the endocytic uptake of Tf647 back to the cell surface, as measured by FACS (Sup Fig 2C). Overall results show that PA does not affect endocytic uptake or recycling of cargo, however, it does impact intracellular localization of endocytic vesicles, which may be crucial for entry of certain viruses.

**Figure 6.**
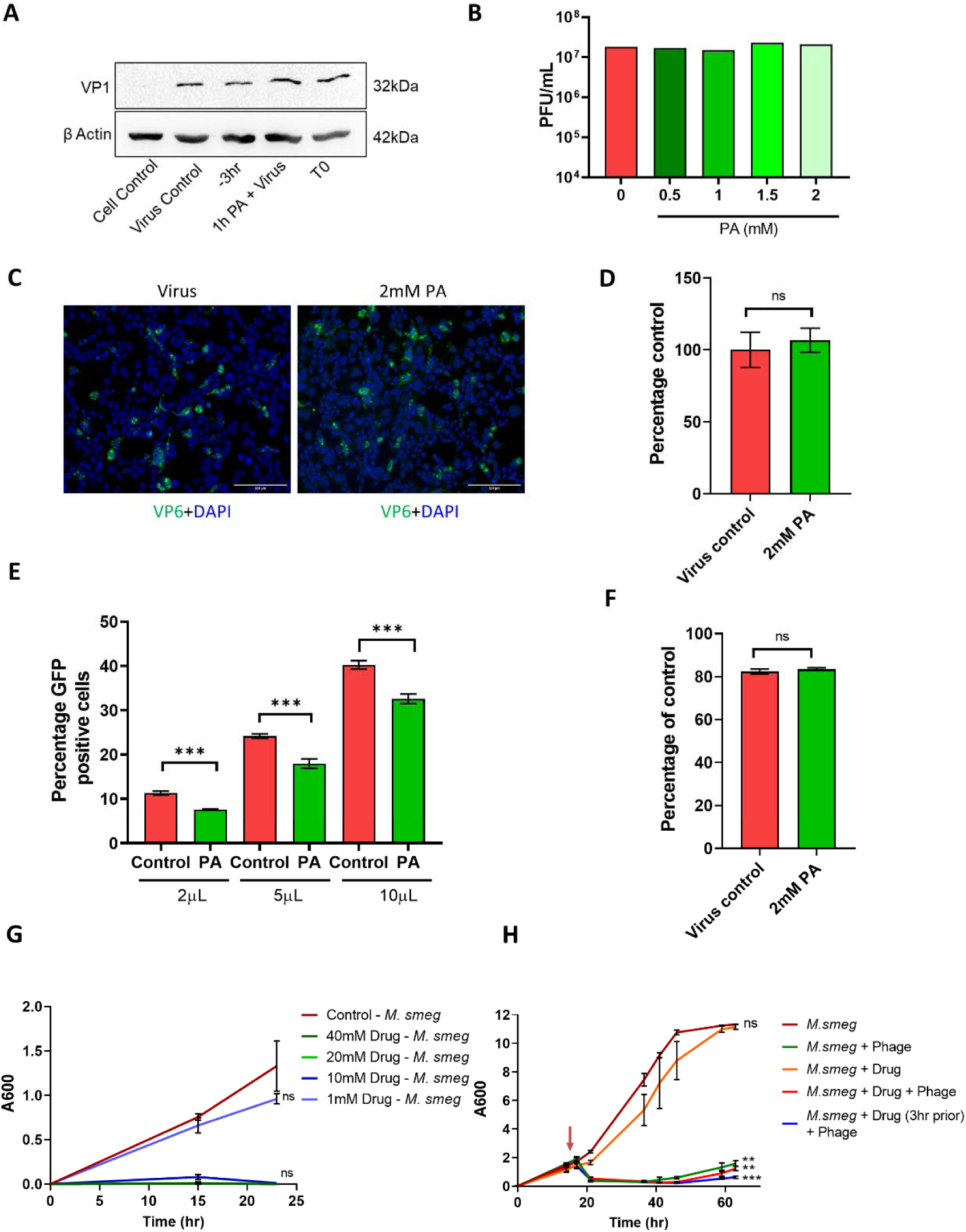
Picolinic acid does not inhibit infection by non-enveloped viruses and bacteriophages. HeLa **c**ells were either pre-treated for 3hr with 2mM PA, infected with 10 MOI CVB3 in presence of the drug and collected 3hr later (−3hr); or treated during infection (T0), or virus and drug were incubated together for 1hr and used for infection (1hr virus+PA). (A) shows quantification of virus infection by western blot using VP1 antibody. (B) HeLa cells were pre-treated with increasing concentrations of PA as indicated and infected with 0.1 MOI CVB3 in the presence of drugs. 48hr p.i, cell culture supernatants were used to quantify infectious virus by plaque assay. (C) HEK293T cells pre-treated with 2mM PA were infected with RRV, fixed with 4% formalin after 12hr and immunolabelled with VP6 antibody and DAPI to label the virus particles and nuclei respectively. (D) Percentage positive cells were quantified using ImageJ/Fiji. (E) HEK293T cells pre-treated with 2mM PA were infected with AAV6-GFP particles in presence of the drug at different volumes as indicated. 48hr p.i, cells were harvested and analyzed for GFP positive cells by flow cytometry. (F) HEK293 cells pre-treated with 2mM PA were infected with 10MOI Adenovirus 5 expressing eGFP in the presence of the drug and harvested 24hr later for quantification of GFP positive cells by flow cytometry. (G) *M.smegmatis* cells in a 48 well plate were treated with increasing concentrations of PA as indicated and OD600 measurements were taken up to 24hr. (H) log-phase secondary bacterial cultures were treated with 1mM PA at regular time intervals as indicated and infected with 10 MOI TM4 mycobacteriophage. OD600 measurements were taken up to 60hr p.i. **p < 0.01; ***p < 0.001; ns - not significant, using two-tailed unpaired t-test or one-way ANOVA with Dunnett’s multiple comparison test, wherever necessary. Error bars represent mean ± standard deviation.

## Discussion

In the previous century, there were 4 global pandemics, 3 by Influenza A viruses and 1 by HIV (Benatar, 2002, Kilbourne, 2006). In the current century, we have already seen 2 global pandemics, the first one caused by H1N1 ‘Swine Flu’ (Scalera and Mossad, 2009) and the second ongoing COVID-19 pandemic caused by SARS-CoV-2 (Fontanet et al., 2021). In addition, there have been several outbreaks of different scales caused by Dengue (Wilder-Smith and Byass, 2016), Chikungunya (Pialoux et al., 2007), Zika (Chang et al., 2016), Ebola (House, 2014), SARS (Cherry and Krogstad, 2004), MERS (Al-Omari et al., 2019), Nipah (Soman Pillai et al., 2020), and several other viruses. The frequency of virus emergence has increased in the recent past, primarily due to increased human activities leading to animal habitat destruction, and climate change (Robert et al., 2020, Morens et al., 2004). Respiratory viruses like Influenza and Coronaviruses, in particular, have shown significant pandemic potential and are likely to cause more outbreaks in the future (Taubenberger and Morens, 2010, Telenti et al., 2021). Vaccine technologies have seen a major advance, especially during the COVID-19 pandemic, as several effective candidates were developed at a lightning speed (CLEVE, 2021). In contrast, the development of antivirals, in general, has been lagging. Antivirals targeting viral factors are difficult to develop and are prone to becoming ineffective due to the rapid emergence of drug-resistant mutants (Debing et al., 2015). Targeting host factors or processes commonly used by a wide range of viruses is an attractive avenue for developing broad-spectrum antivirals. Such antivirals will be well suited for a pandemic scenario, as they are less likely to be affected by viral resistance and will be effective against rapidly emerging variants of the pandemic virus (Adalja and Inglesby, 2019). In general, the viral replication cycle involves hsvsteps of entry, transcription, translation, replication, assembly, and release events. Inhibition of viral entry is the most effective way to prevent viral infection, which is also the mechanism of action of neutralizing antibodies elicited as part of natural or vaccine-mediated immunity (Reading and Dimmock, 2007). Human viruses primarily use either receptor-mediated endocytosis or viral-cellular membrane fusion, or both to enter target host cells (Marsh and Helenius, 2006). The cellular factors involved in these processes are attractive targets to develop broad-spectrum viral entry inhibitors (Mazzon and Marsh, 2019a).

Picolinic acid is a natural metabolite, produced during catabolism of the essential amino acid tryptophan in humans and other mammals through the Kyneurine Pathway (González Esquivel et al., 2017). It is a bidentate chelating ligand, which in humans, is known to aid in the absorption of zinc and other trace elements like iron and chromium from intestines (Evans and Johnson, 1980, Aggett et al., 1989). It is present at varying concentrations in human milk, pancreas, serum, and cerebrospinal fluid (Grant et al., 2009). PA has been reported to exhibit antiviral activity against HIV-1, HSV-2 (Fernandez-Pol et al., 2001), and Chikungunya virus (Sharma et al., 2016), although the mechanism of action remained unclear. In a recent study, PA was shown to interact with cellular E3 Ligase UBR4 and interfere with endocytic vesicle maturation (Kim et al., 2018). Endocytosis is a commonly used mode of viral entry, and UBR4 is required for viral replication of IAV and Dengue viruses, although for different processes (Tripathi et al., 2015, Morrison et al., 2013). Based on this background, we hypothesized that PA may have broad-spectrum antiviral activity, potentially involving inhibition of receptor-mediated endocytosis during viral entry. To test this, we first explored the effects of PA against IAV and found it to be effective against PR8 lab strain and 2009 H1N1 ‘Swine Flu’ strain *in vitro*. Subsequently, we tested PA against SARS-CoV-2 and found the drug to be effective across different VOCs in cell lines of human and non-human primate origin. On further exploration, we observed that PA has an antiviral effect *in vitro*, against Flaviviruses (Dengue, Zika, JEV), HPIV-3, and HSV-1. All these viruses have different genomic organizations and modes of replication; however, a common feature is the presence of the host-derived viral membrane. Also, IAV and Flaviviruses require receptor-mediated endocytosis, but HSV-1 and SARS-CoV-2 can enter cells via direct viral-cellular membrane fusion (Shang et al., 2020). This suggested that PA action may be directed on the viral membrane.

To investigate the MoA of PA, time of addition studies were conducted using SARS-CoV-2 and IAV. Results showed that PA exerts antiviral effect only when added before viral infection, confirming viral entry as its target. For IAV, PA inhibited fusion between viral and cellular endocytic membrane. These data established the inhibitory effect of PA on membrane fusion events during viral entry. Physical examination of IAV by TEM revealed that PA treatment causes disruption in viral membrane integrity, which may be responsible for impaired viral-cellular membrane fusion. It must be noted that PA may have a similar effect on the cellular membrane, however, they can self-repair, but the viral membrane cannot recover from PA-induced damage. Antivirals based on similar principles have been developed before (Wolf et al., 2010). In the case of SARS-CoV-2, in addition to inhibiting viral entry, PA also inhibited Spike-mediated syncytia formation, which involves cell-cell membrane fusion. Notably, we observed more potent *in vitro* antiviral action of PA against SARS-CoV-2, as compared to IAV. It is likely that exogenously added PA will be more effective against more accessible viral-cell membrane fusion (during SARS-CoV-2 entry), as compared to less accessible viral-endocytic membrane fusion (during IAV entry) site. The inhibition of syncytia formation by SARS-CoV-2 may also contribute to enhanced antiviral action of PA against SARS-CoV-2, as compared to IAV. Taken together, these data explain why PA exhibits broad-spectrum *in vitro* antiviral activity against enveloped viruses, even those which are not obligated to endocytosis but do require viral-cellular membrane fusion (for example HSV-1, SARS-CoV-2). Nevertheless, non-enveloped viruses also often use receptor-mediated endocytosis for entry, and hence should be impacted by PA treatment, however, our experiments revealed no effect on the entry of Coxsackie Virus, Rotavirus, and Adenovirus 5. We saw a limited antiviral effect of PA on AAV-6 infection, which may be associated with its effect on cellular endocytosis. Notably, in our experiments PA exhibited no effect on Transferrin uptake or recycling, however, did dysregulate localization of endocytic vesicles, which is consistent with the published report (Kim et al., 2018). Overall, these results indicate that PA primarily acts by interfering with membrane fusion during viral entry (on the cell surface or endocytic compartment), or syncytia formation, however, the role of PA-mediated endocytosis dysregulation is limited in its antiviral action. Nevertheless, membrane fusion and endocytosis are features of viral entry into eukaryotic cells. As expected, PA did not exhibit any antiviral effect on bacteriophage infection in Mycobacteria. This suggests that antiviral action of PA may have evolved in higher organisms, primarily against enveloped viruses, which enter the cell via direct viral-cellular membrane fusion. Recent reports suggest upregulation of the tryptophan metabolic pathway during virus infection (Mehraj and Routy, 2015, Kaur et al., 2021). Also, some of these genes involved in this pathway are potentially upregulated by interferons (Hassanain et al., 1993, Robinson et al., 2003). Taken together, upregulation of picolinic acid can be a natural antiviral response, which needs to be explored further.

Furthermore, we also examined *in vivo* antiviral efficacy of PA against IAV in the murine model and SARS-CoV-2 in the Syrian golden hamster model. We administered PA via IP and oral routes as a prophylactic or therapeutic agent, and it exhibited antiviral activity in all formats. The best outcome was observed when PA was administered prophylactically. This was consistent with its mechanism of action which is inhibition of early events of viral infection. In the case of SARS-CoV-2, inhibition of syncytia formation may also contribute to its *in vivo* antiviral action. PA has been shown to exhibit synergistic activity with IFN-gamma, in promoting the induction of cytotoxic macrophages in mice (Varesio et al., 1990, Ruffmann et al., 1984). Thus, there might be an additional mechanism through which PA may help in resolving a viral infection *in vivo*. PA was better tolerated in Oral administration, and although we tested 20 mg/kg body weight dose, even 100 mg/kg body weight dose was non-toxic in our study and in another study (Ruffmann et al., 1987) which is equivalent to 8 mg/kg in humans (conversions as per https://www.fda.gov/media/72309/download). These data qualify PA as a clinically relevant broad-spectrum antiviral that can be administered orally or through IP route to combat pandemic Influenza and Coronaviruses.

In conclusion, we demonstrate broad-spectrum antiviral action of Picolinic Acid against a wide range of enveloped viruses, including major human viral pathogens, especially pandemic SARS-CoV-2 and Influenza A viruses. We also delineate the mechanism of action of PA and show that it acts by interfering with viral-cellular membrane fusion, virus-induced syncytia formation, and cellular endocytosis. We also provide evidence of excellent tolerance and promising preclinical efficacy of PA against SARS-CoV-2 and Influenza A virus, both through IP and Oral routes of administration. This study paves the way for further development of Picolinic acid as a broad-spectrum antiviral for clinical use and reinforces the utility of host-directed antivirals in combating a wide range of human viruses, especially in a pandemic scenario.

## Limitations of this study

Although the effect of PA on viral membrane has been demonstrated, its effect on cellular membrane has not been examined. Nevertheless, the dose of PA used here was non-toxic to cells, and endocytic uptake and recycling of transferrin were not affected, which rules out potentially adverse effects on cellular membrane. Finally, although PA is a natural metabolite, in this study we have used it as an exogenous antiviral agent. To understand the antiviral role of PA which is produced naturally, genetic, and metabolic modulation of PA production and the subsequent effect on antiviral immunity needs to be investigated in detail using *in vitro* and *in vivo* models.

## Acknowledgments

This work is supported by funding from DBT-Wellcome Trust India Alliance (IA/I/18/1/503613) and DBT-BIRAC (BT/CS0007/CS/02/20) to ST Lab. We acknowledge Infrastructure support from the Crypto Relief Fund, L & T Trust, DST-FIST program, and DBT-IISc partnership program (Phase II) to IISc. RN is supported by the DBT-BIRAC grant to ST Lab. AB is supported by Kishore Vaigyanik Protsahan Yojna (KVPY) fellowship. OK, MS, RY acknowledges support by the MHRD-IISc fellowship. Bacteriophage work in the RA laboratory was supported by Ramanujan Fellowship (SB/S2/RJN-037/2017). We thank Sushma Krishnan, Electron Microscopy Facility, Division of Biological Sciences, IISc, and for assisting TEM imaging. We are thankful to Prof. Adolfo Garcia-Sastre, Prof. Mathew Evans, Prof. Ben Hur Lee (Microbiology, Icahn School of medicine at Mount Sinai, NY, USA), Prof. Andrea Gamarnik, (Fundación Instituto Leloir-CONICET, Buenos Aires, Argentina), and Prof. David Leib (Geisel School of Medicine at Dartmouth, NH, USA) for providing different virus strains.

## Author Contributions

Conceptualization: ST; Methodology: RN, RR, AB, OK, MS, RY, SJ, PRS; Data curation and Analysis: RN, RR, AB, SJ, PRS; Resources: ST, RA, VS, SD, CDR; Validation, Visualization: RN, AB; Writing, Review, Editing: RN, ST, AB, VS, SD, CDR, RA; Funding acquisition, Project administration, Supervision: ST

## EXPERIMENTAL MODEL AND SUBJECT DETAILS

### Ethics Statement

This study was conducted in compliance with institutional biosafety guidelines, (IBSC/IISc/ST/17/2020; IBSC/IISc/ST/18/2021), following the Indian Council of Medical Research and Department of Biotechnology recommendations. All experiments involving animals were reviewed and approved by the Institutional Animal Ethics Committee (Ref: IAEC/IISc/ST/784/2020) at the Indian Institute of Science and conducted in Viral Biosafety level-3 facility. The experiments were performed according to CPCSEA (The Committee for Control and Supervision of Experiments on Animals) guidelines.

### Animal Models

For IAV infection experiments, 4-6 weeks old female BALB/c mice (Central Animal Facility, Indian Institute of Science, Bengaluru, India) were used. Animal experiments involving SARS-CoV-2 infection were performed on 10–12 week-old mixed-gender Syrian golden hamsters (Biogen Laboratory Animal Facility Bengaluru, India) with male and female hamsters housed separately. All animals were housed in groups of four in individually ventilated cages maintained at 23±1 °C temperature and 50±5% relative humidity, given access to standard pellet feed and water ad libitum, and maintained on a 12-hour day/night light cycle at the Viral Biosafety level-3 facility, Indian Institute of Science. All animals were monitored daily during the experiment. An overdose of Ketamine (Bharat Parenterals Limited) and Xylazine (Indian Immunologicals Ltd) was used to sacrifice animals upon completion of the experiment.

### Cell Lines

The following cell lines were used in this study: HEK293T cells expressing human ACE2 (NR-52511, BEI Resources, NIAID, NIH); HEK293T (NCCS, Pune, India), Vero E6 (CRL-1586, ATCC®); Madin-Darby Canine Kidney (NCCS, Pune, India); A549 (NCCS, Pune, India), Calu-3 (ATCC HTB-55), HeLa (ATCC CCL-2) All cell lines were cultured in complete Dulbecco’s modified Eagle medium (12100-038, Gibco) with 10% HI-FBS (16140-071, Gibco), 100 IU/ml Penicillin and 100 μg/ml Streptomycin (15140122, Gibco) supplemented with GlutaMAX™ (35050-061, Gibco).

### Virus Stock and Propagation

The following SARS-CoV-2 isolates were procured from BEI Resources, NIAID, NIH: Isolate Hong Kong/VM20001061/2020, NR-52282; Isolate hCoV-19/England/204820464/2020 (Lineage B.1.1.7; Alpha variant), NR-54000; Isolate hCoV-19/USA/MD-HP01542/2021 (Lineage B.1.351 South Africa; Beta variant), NR-55282; Isolate hCoV-19/USA/PHC658/2021 (Lineage B.1.617.2; Delta Variant), NR-55611; Isolate hCoV-19/Japan/TY7-503/2021 (Brazil P.1 Gamma variant), NR-54982. All these viruses were propagated and titrated by plaque assay in Vero E6 cells as described before (Case et al., 2020). Influenza A virus (IAV) strains namely A/Puerto Rico/8/1934 H1N1 (PR8) and A/California/04/2009 H1N1 (Cal/09) were propagated in 11-day old embryonated chicken eggs and titrated by plaque assay in MDCK cells (Gaush and Smith, 1968). Japanese Encephalitis Virus (JEV) clinical strain P20778 was propagated and titrated in BHK-21 cells. The reporter viruses used in this study include IAV expressing Gaussia luciferase (NS1 Luc) (Tripathi et al., 2015); Dengue virus (DENV Luc), Zika Luc, Human parainfluenza virus (HPIV-3 Luc), and Herpes Simplex Virus (HSV-1 Luc) expressing renilla luciferase, were provided from different laboratories (details in resource table). Adenovirus Serotype 5, Clone Ad5-CMV-hACE2/RSV-eGFP, Recombinant Expressing Human ACE2 was procured from BEI resources (Catalog No. NR-52390). Coxsackie virus B3 (CVB3) was a kind gift from kind gift from Prof. Frank van Kuppeveld. Rhesus monkey rotavirus (RRV) strain was a kind gift from Prof. Durga Rao. C, SRM University, Andhra Pradesh. TM4 mycobacteriophage (Bajpai et al., 2018) was amplified in *M. smegmatis* and phage enumeration was done using the soft agar overlay technique as reported by Kalapala *et al* (Kalapala et al., 2020).

### Bacterial Strains

Primary *Mycobacterium smegmatis* (mc^2^ 155) (a kind gift from Prof. Deepak Saini, Indian Institute of Science) was grown in Middlebrook 7H9 broth (Merck, M0178) supplemented with Glycerol (Fisher scientific, Q24505), ADC (HiMedia, FD019), and 0.1% v/v Tween 80 (Fisher Scientific, YBP338500). A log phase primary culture was inoculated into a secondary culture without Tween 80 and supplemented with 2 mM CaCl_2_ (Fisher scientific, Q12135) to promote efficient infection of phages.

## METHODS DETAIL

### SARS-CoV-2 multicycle infection

HEK293T-ACE2 or Vero E6 cells were seeded in 24-well cell culture plates to reach 70-80% confluency the next day. Cells were pre-treated with 0.25, 0.5, 1, and 2 mM PA for 3hr and infected with 0.1 MOI or 0.001 MOI SARS-CoV-2 Hong Kong in HEK293T-ACE2 and Vero E6 cells respectively in the presence of the drug. For infection, cells were first incubated with 100 µL per well of inoculum and after 1hr adsorption, topped up with 400 µL medium. DMEM containing 2% FBS was used for infection in Vero E6 cells and complete DMEM was used for HEK293T-ACE2 cells. After 48hr, total RNA from infected cells was extracted using TRIzol (Thermo Fisher, 15596018) and viral copy number was estimated by qRT PCR. Cell viability of uninfected, drug-treated cells was measured using Alamar blue cytotoxicity assay (Invitrogen, DAL 1025,) as per the manufacturer’s instructions. Similarly, the effect of PA against SARS-CoV-2 variants of concern was tested by first pre-treating HEK293T-ACE2 and Vero E6 cells with 2 mM PA. Four different variants of concern namely Alpha, Beta, Gamma, or Delta were used to infect HEK293T-ACE2 (0.1 MOI) and Vero E6 (0.001 MOI) cells as mentioned above. After 48hr infection, the total viral RNA copy number from infected cells was estimated by qRT PCR. Vero E6, Calu-3, and HEK293T-ACE2 cells were seeded in 24-well cell culture dishes to reach 80% confluency post 24hr. Cells were then pre-treated with 2 mM PA in triplicates for 3hr and infected with 100 µL per well 0.1 MOI SARS-CoV-2 Hong Kong diluted in complete DMEM in the presence of the drug. After 1hr adsorption, media in wells were topped up with 400 µL DMEM containing 2 mM PA. Viral copy number from infected cells was estimated by qRT PCR 48hr post-infection.

### Quantification of viral load by qRT-PCR

Cells were harvested in TRIzol as per the manufacturer’s instruction. An equal amount of RNA was used to determine viral load using the AgPath-ID™ One-Step RT-PCR kit (Applied Biosystems, AM1005). The following primers and probes targeting the SARS-CoV-2 N-1 gene were used for amplification. Forward primer: 5’GACCCCAAAATCAGCGAAAT3’ and Reverse primer: 5’ TCTGGTTACTGCCAGTTGAATCTG3’, Probe: (6-FAM / BHQ-1) ACCCCGCATTACGTTTGGTGGACC). The Ct values were used to determine viral copy numbers by generating a standard curve using the SARS-CoV-2 genomic RNA standard.

### IAV multicycle infection

MDCK cells were seeded in 24-well cell culture dishes to reach 80-90% confluency after 24hr. Cells were treated with 0.25, 0.5, 1, and 2 mM PA for 3hr, infected with 100 µL per well 0.01 MOI of PR8 in Opti-MEM reduced serum media (Gibco, 31985088) containing 1 µg/mL L-tosylamide-2-phenyl ethyl chloromethyl ketone (TPCK trypsin) (Sigma Aldrich, T1426) in triplicates. For, Cal/09 virus, a single dose of 2 mM PA was used for treatment. Post 1hr virus adsorption, media was topped up to 500 µL. After 48hr, supernatants from each condition were pooled and centrifuged at 2000xg to remove cell debris, before using for plaque assay. In all cases, PA was present in media throughout the experiment. Uninfected cells treated with the different doses of PA, were used for estimation of cell viability by Alamar blue assay.

### IAV plaque assay

MDCK cells were seeded in 12-well plates to reach complete confluency after 24hr. Dilutions (10-fold) of supernatants were prepared in Opti-MEM and 150 µL per well was used to infect cells for 1hr at 37°C with regular shaking every 10 min. The virus inoculum was then removed, and cells were overlaid with 1 mL per well MEM containing 0.6% oxoid agar (Thermo Scientific, LP0028), 1µg/mL TPCK trypsin, and 0.01% DEAE-Dextran hydrochloride (Sigma Aldrich, D9885) and 0.5% Sodium bicarbonate (MP Biomedicals, 194553). Cells were collected 48hr later, fixed with 5% formalin and plaques visualized by crystal violet staining.

### Reporter virus infection

A549 cells at 80-90% confluency in a 24-well dish were treated with 2 mM PA for 3hr in triplicates. Cells were then washed and infected with 100 µL per well DMEM containing 1 µL of DENV Luc or ZIKV Luc; 0.2 µL HPIV-3 Luc or 1 µL HSV-1 Luc. Drug (2 mM PA) was present in the medium throughout the experiment. After 48hr, cells were harvested for detection of firefly and renilla luciferase expression using Dual-Luciferase Reporter Assay System (Promega, E1980) as per manufacturer’s instructions. Luminescence measurements were taken using a TECAN Infinite 200-PRO multiplex reader.

### Flavivirus infection

Confluent A549 cells in a 24-well cell culture plate were pre-treated for 3hr with 2 mM PA and infected with 100 µL per well DMEM containing 0.1 MOI JEV clinical strain P20778 or ZIKV Cambodia. After 1hr adsorption, the wells were topped up with 400 µL DMEM. Drug (2 mM PA) was present in the media throughout the experiment. Cells were then washed with PBS and harvested for western blot analysis 48hr post-infection. The separated proteins were transferred onto the PVDF membrane and probed using mouse anti-Flavivirus envelope 4G2 primary antibody and anti-mouse-HRP conjugated secondary antibody. Actin labeling using Mouse mAb to beta Actin-HRP (Abcam, ab49900) was used as a loading control.

### SARS-CoV-2 infection in hamsters

Experimentally, hamsters under intraperitoneal (IP) Ketamine (150mg/kg) (Bharat Parenterals Limited) and Xylazine (10mg/kg) (21, Indian Immunologicals Ltd) anesthesia were challenged intranasally with 10^5^ plaque-forming units (PFU) SARS-CoV-2 in 100 µL PBS. Two dosage regimens and routes of drug administration were followed for the treatment of animals. The prophylactic treatment used administration of 20 mg/kg/day PA via oral or IP route during -3, -2, and -1 day before infection and therapeutic dosage (oral/IP) involved administering 20 mg/kg/day PA during 1, 2 and 3-day post-infection (dpi). This corresponds to a human equivalent dose of 20 mg/kg (Hamster) x 0.13 (conversion factor) = 2.6 mg/kg (conversions as per https://www.fda.gov/media/72309/download). A total volume of 200 µL PA dissolved in PBS was used for both oral and IP routes of administration. The total bodyweight of animals was recorded every day until the end of the experiment at 4 dpi when the animals were sacrificed. Total lungs were harvested, weighed, and processed for histopathological analysis. One portion was used for RNA extraction using TRIzol and subsequent viral RNA copy number estimation by qRT-PCR as described previously.

### Influenza virus infection studies in mice model

To assess toxicity, animals were treated with 20 or 100 mg/kg PA by either intraperitoneal (IP) or oral routes. One group served as PBS (untreated) control. The body weight and general health of animals were measured every day for up to 9 days post-treatment. Infection/treatment groups were divided into two, one group receiving 20 mg/kg PA prophylactically and the other, therapeutically. This corresponds to a human equivalent dose of 20 mg/kg (Mouse) x 0.08 (conversion factor) = 1.6 mg/kg (conversions as per https://www.fda.gov/media/72309/download). For infection, mice under intraperitoneal (IP) anesthesia with Ketamine (90 mg/kg) (Bharat Parenterals Limited) and Xylazine (4.5 mg/kg) (21, Indian Immunologicals Ltd) were challenged intranasally with 50 PFU of PR8 WT virus in 40 µL PBS. Two dosage regimens and routes of drug administration were followed for the treatment of animals. The prophylactic treatment used administration of 20 mg/kg/day PA via oral or IP route for 6hr before infection and therapeutic dosage (oral/IP) involved administering the same amount of drug 3hr p.i. One-half of the animals were sacrificed at 4 dpi and lungs were collected for plaque assay quantification of IAV and histology analysis. For remaining animals, total bodyweight and survival were recorded until the end of the experiment at 7 dpi. For plaque assay, lung samples were collected in DMEM containing 0.3% BSA, homogenized, and centrifuged at 5000xg for 10 min at 4°C to pellet tissue debris. The supernatant was used for plaque assay as mentioned previously.

### Histopathology

Lung tissue samples were fixed in 10% buffered formalin, embedded in paraffin blocks and tissue sections of 4-6 μm thickness made using a microtome. The sections were then stained with Hematoxylin and Eosin and examined by light microscopy as previously described (Chan et al., 2020). Clinical scoring for hamster lung samples was done based on three different criteria namely: Alveolar edema; vascular and perivascular infiltration; alveolar thickening and infiltration. In the case of mice, three different clinical criteria were observed, namely: vascular infiltration, alveolar infiltration, and interstitial pneumonia. In both cases, scoring was done based on the severity on a scale of 1-4 (1-mild, 2-moderate, 3-severe, 4-very severe).

### Immunofluorescence assay

Cells on glass coverslips were washed once with PBS, fixed with 4% PFA for 10 min, and permeabilized for 10 min with PBS containing 1% Tween-20. Cells were then washed and incubated in blocking buffer (PBS with 0.3% Tween 20, 2% BSA) for 1hr. Overnight incubation with primary antibody diluted in blocking buffer was followed by washing and incubation with secondary antibody in blocking buffer containing 0.1 µg/mL DAPI (Sigma Aldrich, D9542). Finally, cells were incubated with PBS containing 50 mM ammonium chloride (Fisher Scientific, 21405) for de-quenching, washed, and the cells on coverslips were mounted on glass slides using ProLong™ Diamond Antifade Mountant (Invitrogen, P36961).

### Western Blot

Cells were washed with 1x PBS (162528, MP Biomedicals), lysed with 1x Laemmli buffer (1610747, BIO-RAD), and heated at 95°C for 10 min. Cell lysates were then subjected to standard SDS-PAGE and separated proteins were transferred onto a PVDF membrane (IPVH00010, Immobilon-P; Merck). The membrane was incubated in a blocking buffer containing 5% Skimmed milk (Sigma Aldrich, 70166) in 1xPBS containing 0.05% Tween 20 (Sigma-Aldrich P1379) (1xPBST) for 2hr with slow rocking at room temperature (RT). Primary antibody incubation in blocking buffer was done for 14hr at 4°C with gentle rocking, after which the membrane was washed with 1x PBST and incubated for 2hr with secondary antibody in blocking buffer at RT. After a further wash with 1x PBST, the blots were developed using Clarity Western ECL Substrate (Bio-Rad, 1705061).

### IAV time of addition assay

A549 cells were seeded in 24-well cell culture plates containing glass coverslips to reach 70-80% confluency the next day. For infection, cells were washed once with warm PBS and incubated with 100 µL per well PR8 WT virus (2 MOI) diluted in OptiMEM. The plates were rocked every 10 min to ensure even distribution of inoculum. After 1hr adsorption, the medium in wells was topped up with 400 µL OptiMEM. Effects of PA on early and late events during IAV infection were studied at 3 and 9hr time points post-infection, respectively. In the former, cells were pre-treated with 2 mM PA for 3hr, infected, and collected at 3hr post-infection. Alternatively, PA treatment was done only at 6hr post-infection and cells were harvested after a further 3hr incubation period. The direct effect of the drug on virus particles was studied by incubating virus inoculum in Opti-MEM containing 2 mM PA at 37°C for 60 min, followed by which the virus-drug mixture was used to infect cells as mentioned above. Post 1hr adsorption, the medium was topped up with 400 µL Opti-MEM without drug. In all cases, cells were harvested for both IFA and western blot analysis. For IFA, Anti-mouse Influenza virus NP (HT103) was used as the primary antibody. Goat anti-Mouse IgG (H+L) Cross-Adsorbed Secondary Antibody, Alexa Fluor 488 (Invitrogen, A11001) was used as the secondary antibody. Images were acquired using an EVOS M5000 Imaging system. Quantification of NP positive cells relative to the total number of DAPI positive cells in 5 different fields was performed using ImageJ/Fiji software. Primary antibody incubation for western blot was done with Anti-mouse Influenza virus NP (HT103) and secondary antibody with Goat Anti-Mouse IgG - H&L Polyclonal Antibody, HRP conjugated (Abcam, ab6789).

### SARS-CoV-2 time of addition assay

Vero E6 and HEK293T-ACE2 cells were seeded in 24-well cell culture plates containing glass coverslips to reach 70-80% confluency the next day. For infection, cells were washed once with warm PBS and incubated with 100µL per well 10 MOI SARS-CoV-2 diluted in complete DMEM. The plates were rocked every 10 min to ensure even distribution of inoculum. After 1hr adsorption, the media in wells were topped up with 400µL OptiMEM. Effects of PA on early and late events during SARS-CoV-2 infection were studied at 3 and 9hr time points p.i, respectively. In the former, cells were pre-treated with 2 mM PA for 3hr, infected, and collected at 3hr post-infection. Alternatively, cells were infected and treated simultaneously (T0) and collected after 3hr. Direct effects of PA on virus particles were tested by incubating the virus with 2 mM PA for 1hr at 37°C (1hr virus+PA), which was then used for infection. No additional drug was added at the time of infection. The effect of PA on late events was studied by treating cells only during 6hr post-infection, cells were then harvested after a further 3hr incubation. Once added, PA was present in the medium throughout the remaining duration of the experiment. In all cases, cells were harvested for both IFA and western blot analysis. The antibodies used for IFA include SARS-CoV-2 spike primary antibody (GTX632604, GeneTex) and goat anti-Mouse IgG (H+L) Cross-Adsorbed Secondary Antibody, Alexa Fluor 488. Quantification of spike-positive cells was done using ImageJ/Fiji. Western blot analysis of viral proteins used Polyclonal Anti-SARS-Related Coronavirus 2 Spike Glycoprotein (BEI, NR-52947) and Goat Anti-Rabbit IgG - H&L Polyclonal antibody, HRP Conjugated (Abcam, ab6721).

### Pseudotyped SARS-CoV-2 particle production and transduction

Pseudotyped particles bearing the SARS-CoV-2 spike protein were produced as reported before (Hashimoto et al., 2017). Briefly, HEK293 293T cells were seeded in 10 cm cell culture dishes to reach 50-60% confluency post 24hr and transfected with 2.5µg each of the following plasmids: Vector pHDM Containing the SARS-Related Coronavirus 2, Wuhan-Hu-1 Spike Glycoprotein (BEI, NR-52514); Lentiviral Backbone, Luc2; ZsGreen (BEI, NR-52516); Helper plasmid, Gag; pol (BEI, NR-52517); Helper plasmid, Tat1b (BEI, NR-52518) and Helper plasmid, Rev1b (BEI, NR-52519) using Lipofectamine-2000 transfection agent (Invitrogen, 11668019). The supernatants were pooled together 60hr post-transfection, centrifuged at 5000xg for 10 min at 4°C to remove cell debris, and finally passed through a 0.45µm syringe filter (Sigma Aldrich, 9913-2504) before being used for transduction. HEK293T-ACE2 and Calu-3 cells were seeded in 96-well dishes to reach 60-70% confluency after 24hr. Cells were treated with 2, 1, 0.5, and 0.2mM PA and 3hr later, transduced with 100µL per well pseudotyped SARS-CoV-2 particles containing 5µg/mL polybrene (Merck, TR-1003-G). The different concentrations of PA were present throughout the experiment. Post transduction (60hr), cells were washed once with PBS and processed for detecting luciferase expression using a Firefly luciferase assay kit (Promega, E4550) as per the manufacturer’s instructions. Luminescence measurements were taken using a TECAN Infinite 200-PRO multiplex reader.

### Influenza polymerase Assay

HEK293293T cells were seeded in a poly-L-lysine (Sigma Aldrich, P915) coated 24-well cell culture dish to reach 50-60% confluency the next day. Cells were then co-transfected with plasmids encoding components of the Influenza virus polymerase (P) complex including 50ng each of PA, PB1, PB2; 200ng of Nucleoprotein (NP); 50ng NP-firefly Luciferase; and 100ng pRLTK (mammalian vector for the weak constitutive expression of wild-type Renilla luciferase) as an internal control. Post 3hr transfection, cells treated with 2mM PA, harvested at 8 and 12hr post-treatment for detection of firefly and renilla luciferase expression using Dual-Luciferase Reporter Assay System (E1980, Promega) as per manufacturer’s instructions. Luminescence readings were taken using a TECAN Infinite 200-PRO multiplex reader.

### SARS-CoV-2 spike induced syncytia assay

Vero E6 cells were seeded in a 10 cm cell culture dish to reach 50-60% confluency after 24hr. Cells were transfected with 5 µg plasmid expressing SARS-Related Coronavirus 2, Wuhan-Hu-1 Spike Glycoprotein (BEI resources NR-52514) using Lipofectamine 2000 transfection reagent, as per manufacturer’s instructions. After 24hr, cells were trypsinized and mixed with an equal number of normal un-transfected Vero E6 cells to form a homogenous cell suspension. These cells were then seeded in a 24 well-cell culture dish containing glass coverslips, at a density of 1,00,000 cells per well. After 1hr, cells were treated with 0.25, 0.5, 1 and 2 mM PA in triplicates, and incubated at 37°C, 5% CO2. Non-treated cells and normal un-transfected Vero E6 cells serve as positive and negative controls respectively. After 24hr incubation, the cell culture plate was placed on ice to arrest endocytosis. Cells were then washed once with cold PBS and incubated with 10 µg/mL Wheat Germ Agglutinin (WGA) (Invitrogen, W11261) for 3 min, after which cells were fixed with 4% PFA for 10 min. This was followed by incubation of cells in PBS containing 0.1µg/mL DAPI (Sigma Aldrich, D9542) for 10 min to label nuclei. Finally, cells were washed with PBS and the coverslips were mounted on glass slides using ProLong Diamond Antifade Mountant (Invitrogen, P36970). Cells were imaged using an EVOS M5000 fluorescence microscope and the area of syncytia across different conditions was quantified by drawing ROIs using ImageJ/Fiji.

### Influenza virus membrane fusion assay

#### Virus labeling

IAV particles were labeled using Octadecyl Rhodamine B Chloride (R18) (Invitrogen, O246) as reported earlier, with minor modifications (Hoekstra et al., 1984). A total volume of 1mL PR8 wild type virus (2×10^9^ PFU/mL, titrated by plaque assay) was centrifuged at 2.5xg for 5 min to remove any debris, and placed on ice. R18 dye was added to the virus at 20µM final concentration while simultaneously subjecting to continuous vortexing for 2 min, followed by which the virus-dye mixture was incubated for 60 min on a rocker at RT. After a further 60 min incubation ice, the preparation was centrifuged at 25,000xg for 3hr at 4°C to remove unbound dye. Finally, the concentrated R18 labeled virus preparation was carefully removed from the tube and re-suspended in 200µL NTC buffer (100 mM NaCl, 20 mM Tris-HCl pH 7.4, 5 mM CaCl2).

#### Membrane fusion assay

The methodology for detection of IAV membrane fusion in endosomes was adapted from a previously reported protocol, with few modifications (Hoffmann et al., 2018), Briefly, MDCK cells in suspension were pre-treated for 3hr with 2mM PA, 10µM ammonium chloride (NH4Cl) or 10µM Chloroquine (CQ) at 37°C. Cells were then washed with ice-cold PBS and infected with 10MOI R18 labeled virus in Opti-MEM containing 1µg/mL TPCK trypsin, and placed on ice. Post 1hr adsorption, cells were transferred to a pre-chilled opaque flat-bottom 96-well cell culture plate (136101, Nunc) and placed in a TECAN Infinite 200-PRO multiplex reader pre-set at 37°C. Fluorescence intensity measurements (Ex 560/Em 590) were recorded at every 5 min interval for up to 90 min.

### Characterizing effects of PA on endocytosis

#### Transferrin uptake assay

A549 cells in a 24-well dish were treated or not with 2mM PA for 2hr followed by washing of cells with PBS and starvation for 1hr in the presence of DMEM without FBS for 1hr. Cells were then washed and incubated with 25µg/mL Transferrin Alexa Fluor 647 Conjugate (Invitrogen, T23366) for 1hr. PA was present during the entire course of the experiment. Finally, cells were washed twice with PBS, trypsinized, and re-suspended in PBS containing 0.03% BSA for analysis in a Cytoflex flow cytometer. Results were analyzed using CytExpert software.

Localization of transferrin labeled vesicles: Vero E6 cells were seeded on glass coverslips in a 12 well dish to reach 70-80% confluency the next day. After 3hr treatment with 2mM PA, cells were washed once with ice-cold PBS and the plate was placed on ice for 10 min before incubating with Opti-MEM containing PR8 wild type virus (100 MOI) and 25µg/mL Transferrin 647 (Tf-647) for 1hr on ice. Cells were washed twice with PBS and the plate moved to 37°C. After a 15 min chase, cells were fixed with 4% PFA, labeled with anti-influenza virus HA antibody (PY102) followed by anti-mouse Alexa 488 secondary antibody to label the virus particles. Nuclei were labeled using 0.1µg/mL DAPI. Images were acquired using a Zeiss LSM-880 Multiphoton microscope. The distribution of transferrin labeled vesicles from the perinuclear region to cell periphery was quantified by drawing region of interest (ROIs) up to 20µm away from the nuclei. The fluorescence intensity of Tf-647 labeled vesicles was then measured along this distance using ImageJ/Fiji.

### Adeno Associated Virus - 6 productions and infection

#### AAV6 production

AAV6 particles were produced as per a previously published protocol (Negrini et al., 2020) with few modifications. Briefly, HEK293T cells were seeded in 2 X T75 flasks to reach 50-60% confluency the next day. For each flask, the following plasmids were transfected using Lipofectamine 2000 transfection reagent (Invitrogen, 11668019) as per manufacturer instructions: 17.7 μg pAdDeltaF6(Addgene 112867), 7.9 μg pRepCap6 (Addgene 110770), and 5.9 μg pAAV-CAG-GFP (Addgene 37825). After 60hr, the cells and medium mixture were pooled and transferred to a 50mL conical tube. 3mL Chloroform (Q12305, Qualigens) was added, vortexed gently for 5 min, and 8mL 5M NaCl was added. The tube was then centrifuged for 5 min at 3000 × g, 4°C, and the aqueous phase was transferred to a fresh tube. 10mL of 50% (v/v) PEG 8000 (Sigma, P-2109) was added, vortexed briefly, and incubated for 1hr on ice before centrifuging for 30 min at 3000 × g, 4°C. The supernatant was then discarded, pellet re-suspended in 1.5mL HEPES (H5303, Promega), vortexed for 2 min, and following components were added: 3.5µL of 1M MgCl_2_ (HiMedia, MB237), 14µL DNase I (NEB, M0303S) and 1.4µL of 10μg/μL RNase A (Thermo Scientific, EN0531). The contents were incubated for 20 min at 37°C, equal volume to chloroform added to the tube, and mixed well before centrifuging for 5 min at 3000 × g. The aqueous phase was then transferred to a new tube, followed by which the contents were passed through a 100kDa Amicon Ultra-0.5 Centrifugal Filter Unit (Merck-Millipore, UFC510008) by centrifugation for 5 min at 14,000 × g. The column was washed twice with PBS and AAV particles eluted into a fresh tube by centrifugation at 1000 × g for 2 min.

#### Infection with AAV6 particles

HEK293 293T cells were seeded in a poly-L-lysine coated 24-well dish to reach 60-70% confluency the next day. Cells were pre-treated or not with 2mM PA for 3hr and infected with 100uL complete DMEM per well containing three different volumes i.e 2, 5, and 10uL AAV6 particles. After 1hr, the medium was topped up with 400uL complete DMEM. PA was present in the medium for the entire duration of the experiment. After 48hr, cells were trypsinized and re-suspended in PBS containing 3% FBS (FACS buffer). The number of GFP positive cells was analyzed using a Cytoflex (Beckman Coulter) flow cytometer and results were analyzed using CytExpert software.

#### Adenovirus 5 infection

For infection studies, HEK293 cells were pre-treated for 3hr with 2mM PA and infected with 10 MOI AAV5-eGFP in the presence of the drug. After 24hr, cells were trypsinized, resuspended in FACS buffer, and used to quantify the total number of GFP positive cells by flow cytometry analysis. The drug was present in the treated conditions throughout the experiment.

### Coxsackievirus B3 infection

#### Virus preparation

CVB3 virus was prepared as reported earlier (Klump et al., 1990, van Ooij et al., 2006). Briefly, pCB3/T7 DNA was linearized using SalI-HF enzyme (R3138S, NEB) and CVB3 RNA was produced by *in vitro* transcription reaction. The infectious RNA was transfected in HeLa cells and the cell culture supernatant was harvested after 48hr. The virus was amplified in one passage. For virus titer calculation, plaque assay was performed in Vero E6 cells and plaque-forming units per milliliter were estimated.

#### Plaque assay

Vero E6 cells were seeded in a 12-well plate at a confluency of approximately 90% and infected with serially diluted CVB3-containing cell culture supernatant and incubated at 37℃ for 1hr with gentle swirling of the medium at every 10-15 min interval. After virus adsorption, cells were washed with PBS and overlaid with a 1:1 mixture of 2X DMEM and 1.6% Low melting agarose (Sigma-Aldrich, A9414), and incubated at 37℃. After 48hr, cells were fixed using 4% paraformaldehyde for 1hr and stained with 1 % crystal violet solution.

#### Infection

To study the early effects of PA, HeLa cells were pre-treated for 3hr, infected with 10 MOI CVB3, and collected at 3hr post-infection. Alternatively, cells were infected and treated simultaneously (T0) and collected after 3hr. A mixture of virus and drug incubated for 1hr (1hr PA+Virus) was used for infection to test the effects of PA on virus particles. No additional drug was added here post-infection. Expression levels of VP1 protein were detected by western blot using monoclonal mouse anti-enterovirus primary antibody Clone 5-D8/1 (Dako, M7064,) and anti-mouse antibody-HRP (Sigma Aldrich, A4416).

#### Rotavirus infection

A working concentration of RRV was prepared by diluting the virus stock 2-fold with complete DMEM containing a final concentration of 2µg/mL TPCK trypsin. The mixture was incubated at 37°C for 30 min and further diluted 2-fold in DMEM without serum. This mixture was then used to infect HEK293 cells that were pre-treated for 3hr with 2mM PA. A volume of 100µL per well, in a 24 well plate was used for 1hr adsorption, after which the wells were topped up with 400µL serum-free media containing PA. After 12hr, cells were fixed with 4% formalin and immunolabeled with primary mouse anti VP6, and anti-mouse secondary Alexa Fluor 488 antibodies to detect virus-infected cells by IFA.

#### Drug cytotoxicity assay

A549 cells were seeded in 96-well cell culture dishes and 24hr later treated with 0.02, 0.2, 2, and 20mM PA in triplicates. Cells were then incubated at 37°C, 5% CO2, and cytotoxicity was measured at 12, 24, 36, 48, 60, and 72hr post-treatment using AlamarBlue™ Cell Viability Reagent (Thermo Fisher, DAL 1025) as per manufacturer’s instructions. Similarly, cytotoxicity assay in MDCK cells measured at 48hr post-treatment was done using 0.2, 0.5, 1, and 2 mM PA, in parallel with the multicycle IAV infection experiment mentioned below.

#### Transmission Electron Microscopy

PR8 WT virus of stock titer 2×10^9^ PFU/mL grown in 11 day old embryonated chicken eggs was passed through 0.45µm syringe filter and concentrated by ultracentrifugation at 25,000xg for 2hr at 4°C. The concentrated virus prep was then incubated with either distilled water (vehicle control) or 2 mM PA. Aliquots (2-3 μL) of the virus samples were applied to a Formvar/carbon-covered 300 mesh copper grid (Ted Pella, 01753-F) which was hydrophilized by glow discharging at 8 W for 60 s directly before use. After 2 min, the excess sample was removed using Whatman filter paper (Sigma Aldrich,1001125). 5 µL of negative stain 2% uranyl acetate (SRL Lab, 81405) was added to the grid and incubated for 40 secs, after which excess stain was removed. The negative staining step was repeated 3 times and the grid was air-dried for 10 min before imaging. The grids were imaged using a Talos L120C transmission electron microscope equipped with a LaB_6_ electrode operating at an acceleration voltage of 120 kV. Images of the virus were recorded using a 4k Å∼ 4k Ceta CMOS camera.

### Mycobacteria and TM4 bacteriophage experiments

#### Bacterial toxicity assay

PA stock solution of 1M concentration was prepared in sterile deionized water and diluted to obtain different concentrations (1 mM, 5 mM, 10 mM, 20 mM, and 40 mM), in a 48-well plate. A total of 2 × 10^5^ cells of *M. smegmatis* MC^2^ 155 were added to each of the wells. The 48-well plate was placed in a rotary shaker incubator at 37°C for 24hr. Readings were taken periodically using a Tecan Spark multi-mode plate reader at 600 nm.

#### Effect of drug on TM4 mycobacteriophage infection

To study the effect of the drug on TM4 phage growth and activity, 7H9 broth supplemented with ADC growth supplement (HiMedia, FD019) and Calcium chloride (Fisher Scientific Q12135) was prepared and inoculated with 100 µL of log-phase secondary bacterial culture (OD 1 – 2) per 5mL of the media. 1 mM of the drug was added to culture tubes at the appropriate time (either at the start of the experiment or 3hr before adding the phage in the mid-log phase). The cultures were then incubated at 37°C with rotary shaking at 180 pm. For phage-treated samples, a 10 MOI TM4 mycobacteriophage was added at a specified interval of the mid-log phase. For optical density (OD) measurements, 100 µL of bacterial culture at various time intervals was diluted 10 times in media and pipetted several times to obtain a uniform cell suspension. Readings were taken using Jenway 7205 UV/Visible Spectrophotometer at 600 nm against a media blank.

## GRAPHICAL REPRESENTATIONS AND STATISTICAL ANALYSIS

All statistical analyses were performed using GraphPad Prism 8.4.3 (GraphPad Software, USA). In all cases, p-value < 0.05 is considered significant. Graphical illustrations were created using http://biorender.com/.

---

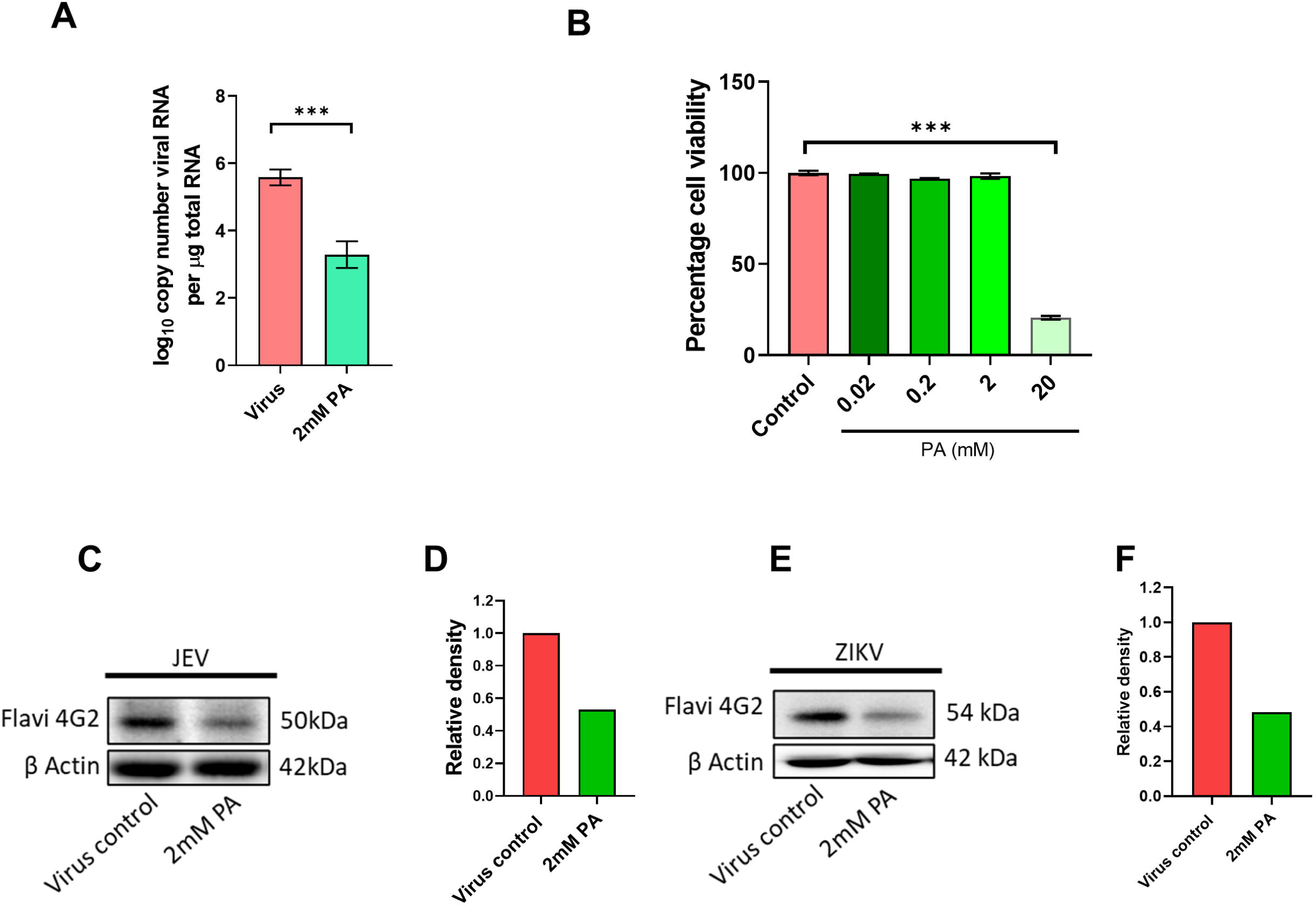

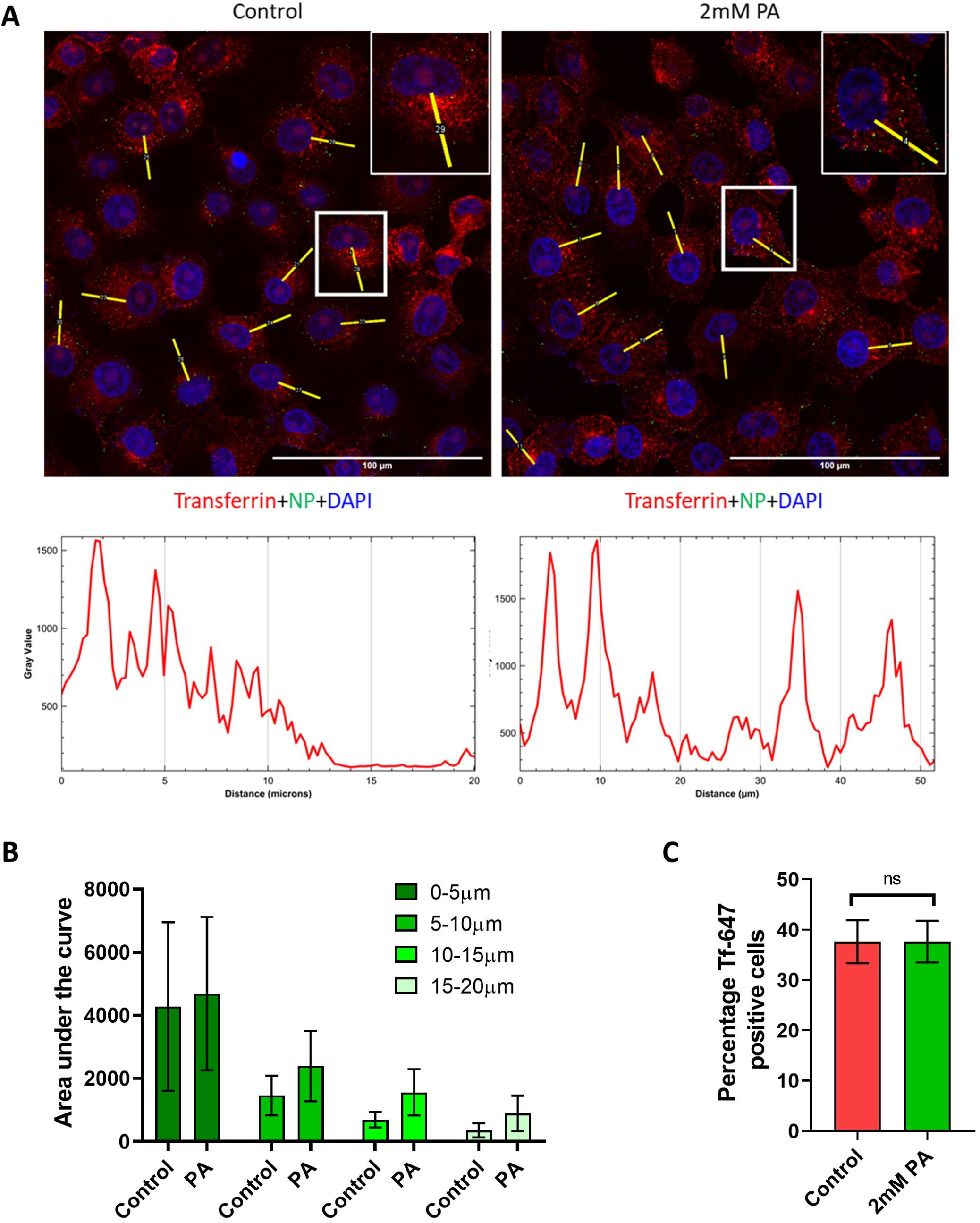

