## Supplementary File for "A natural broad-spectrum inhibitor of enveloped virus entry, effective against SARS-CoV-2 and Influenza A Virus in preclinical animal models"

**Supplementary Information**

**Supplementary figures**

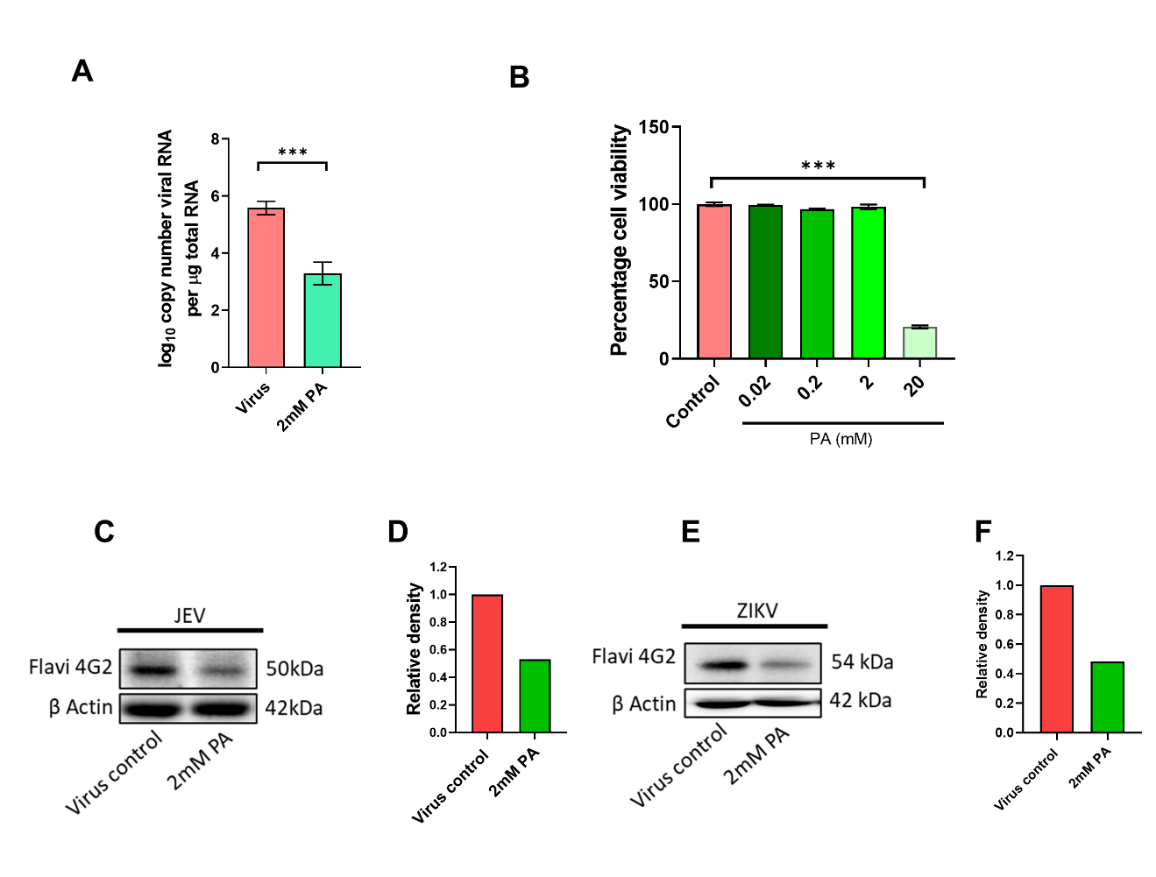
**Supplementary Figure 1.** (A)Calu-3 cells were pre-treated for 3hr with 2mM PA, infected with 0.1 MOI SARS-CoV-2 Hong Kong, and 48hr later, vRNA copy number was estimated by qRT PCR. (B) A549 cells were treated with increasing doses of PA as indicated and cell viability was measured by Alamar Blue assay at 48hr post-treatment. (C-F) A549 cells pre-treated with 2 mM PA for 3hr were infected with 0.1 MOI of either (C,D) JEV or (E,F) ZIKV in the presence of the drug. Cells were collected 48hr p.i and expression of flavivirus envelope protein was detected by western blot. The relative density of bands was analyzed by ImageJ/Fiji (D,F). ***p < 0.001, using two-tailed unpaired t-test or one-way ANOVA with Dunnett’s multiple comparison test, wherever necessary. Error bars represent mean ± standard deviation.

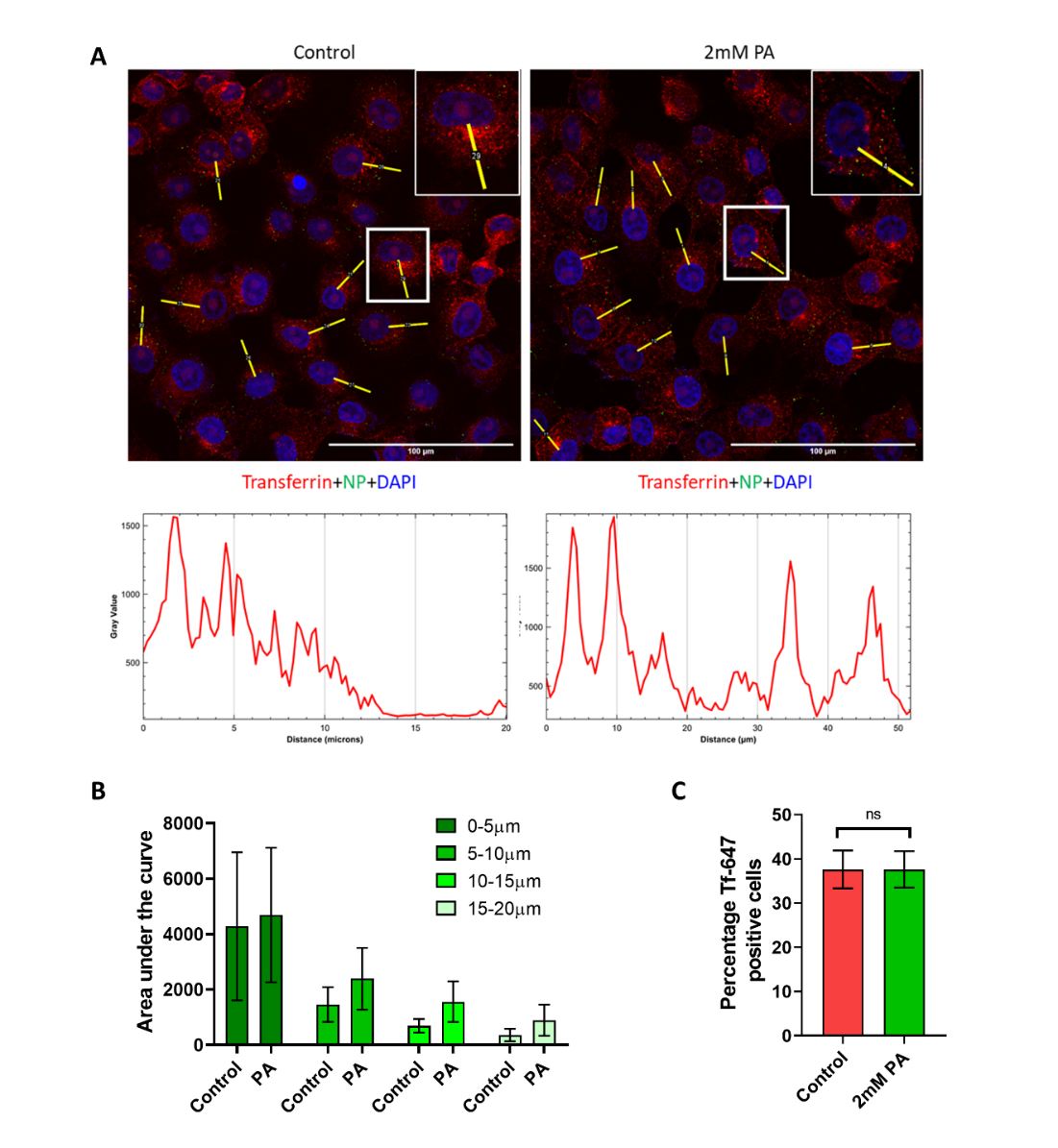
**Supplementary Fig 2: Picolinic Acid has limited effect on Cellular Endocytic Process.** (A-B) PA treated Vero E6 cells were incubated with Tf647 and 100MOI PR8 WT virus for 1hr on ice and moved to 37°C. After 1hr, cells were washed and incubated for 15 min at 37°C before fixation with 4% formalin. (A) Confocal images with nuclei are shown in blue and Tf647 labeled vesicles in red. ROIs were drawn to quantify fluorescence intensities of Tf647 labeled vesicles up to 20µm away from the nuclei using ImageJ/Fiji and the results were plotted as a histogram. (B) shows the area under the curve measured at every 5 µm distance from the nuclei up to 20 µm. (C) A549 cells pre-treated with 2mM PA were pulsed with 25µg/mL Tf647 for 1hr in the presence of PA, washed, and analyzed by flow cytometry to quantify the percentage of Tf647 positive cells.

**Materials Table**

| REAGENT or RESOURCE | SOURCE | IDENTIFIER |
| --- | --- | --- |
| **Antibodies** | | |
| Anti-mouse Influenza virus NP (HT103) | Center for Therapeutic Antibody Development (CTAD), Icahn School of Medicine at Mount Sinai (ISMMS), New York, USA | HT103 |
| Goat anti-Mouse IgG (H+L) Cross-Adsorbed Secondary Antibody, Alexa Fluor 488 | Thermo Fisher Scientific | Cat# A-11001, RRID:AB_2534069 |
| Goat Anti-Mouse IgG - H&L Polyclonal Antibody, HRP Conjugated | Abcam | Cat# ab6789, RRID:AB_955439 |
| Goat Anti-Rabbit IgG - H&L Polyclonal antibody, Hrp Conjugated | Abcam | Cat# ab6721, RRID:AB_955447 |
| SARS-CoV-2 (2019-nCoV) Nucleoprotein / NP Antibody, Rabbit MAb | Sino Biological | Cat# 40143-R019, RRID:AB_2827973 |
| SARS-CoV / SARS-CoV-2 (COVID-19) spike antibody [1A9] | GeneTex | Cat# GTX632604, RRID:AB_2864418 |
| Polyclonal Anti-SARS-Related Coronavirus 2 Spike Glycoprotein (IgG, Rabbit), | BEI Resources, NIAID, NIH | NR-52947 |
| Anti-mouse Flavivirus envelope 4G2 | CTAD, ISMMS, New York, USA | 4G2 |
| Monoclonal Mouse anti-Enterovirus Clone 5-D8/1 | Dako | Cat# M7064 |
| Mouse mAb to beta Actin [AC-15] (HRP) | abcam | Cat#ab49900 |
| Anti-Mouse IgG (whole molecule)–Peroxidase antibody produced in goat | Sigma Aldrich | Cat#A4416 |
| **Bacterial and virus strains** | | |
| *Mycobacterium smegmatis* MC^2^ 155 | Gift from Deepak Saini, Indian Institute of Science | N/A |
| SARS-CoV-2 (Isolate Hong Kong/VM20001061/2020, NIAID, NIH) | BEI Resources, NIAID, NIH | NR-52282 |
| Isolate hCoV-19/Japan/TY7-503/2021 (Brazil P.1) | BEI Resources, NIAID, NIH | NR-54982 |
| Isolate hCoV-19/USA/PHC658/2021 (Lineage B.1.617.2) | BEI Resources, NIAID, NIH | NR-55611 |
| Isolate hCoV-19/England/204820464/2020 (Lineage B.1.1.7) | BEI Resources, NIAID, NIH | NR-54000 |
| Isolate hCoV-19/USA/MD-HP01542/2021 (Lineage B.1.351) | BEI Resources, NIAID, NIH | NR-55282 |
| A/Puerto Rico/8/1934 (PR8) | Kind Gift from Prof. Adolfo Garcia-Sastre (ISMMS, NY) | (Tripathi et al., 2015) |
| A/California/04/2009 H1N1 (Cal/09) | Kind Gift from Prof. Adolfo Garcia-Sastre (ISMMS, NY) | (Tripathi et al., 2015) |
| Viet Nam/1203/04 H5N1 (HALo) | Kind Gift from Prof. Adolfo Garcia-Sastre (ISMMS, NY) | (Tripathi et al., 2015) |
| Japanese Encephalitis Virus clinical strain P20778-GIII | Kind Gift from Prof. Vijaya S. (Microbiology & Cell Biology, Indian Institute of Science, India) | (Krishna et al., 2009) |
| Zika Virus (Uganda 1947 (Strain MR 766; GenBank HQ234498.1) | Kind Gift from Prof. Adolfo Garcia-Sastre (ISMMS, NY) | (Tripathi et al., 2017) |
| IAV expressing Gaussia luciferase (NS1 Luc) | Kind Gift from Prof. Adolfo Garcia-Sastre (ISMMS, NY) | (Tripathi et al., 2017) |
| Dengue virus expressing renilla luciferase (DENV Luc). | Kind gift from Prof. Andrea Gamarnik (Fundación Instituto Leloir-CONICET, Buenos Aires, Argentina), | (Samsa et al., 2012) |
| Zika virus expressing renilla luciferase. | Kind Gift from Prof. Matthew J. Evans (ISMMS, NY) | (Schwarz et al., 2016) |
| Human parainfluenza-3 virus expressing renilla luciferase (HPIV-3 Luc). | Kind Gift from Prof. Benhur Lee (ISMMS, NY) | (Beaty et al., 2017) |
| Herpes Simplex Virus -1 expressing renilla luciferase. | Kind Gift from Prof. David Lieb, Geisel School of Medicine at Dartmouth, NH, USA | (Summers and Leib, 2002) |
| Adenovirus Serotype 5, Clone Ad5-CMV-hACE2/RSV-eGFP, Recombinant Expressing Human ACE2 | BEI Resources, NIAID, NIH | NR-52390 |
| Coxsackie virus B3 | Kind gift from Prof. Frank van Kuppeveld | N/A |
| Rhesus monkey rotavirus (RRV) | Kind gift from Durga Rao, SRM University | (Dhillon and Rao, 2018) |
| TM4 mycobacteriophage | Kind gift from Rachit Agrawal, Indian Institute of Science  Bajpai et al., 2018 | N/A |
| **Chemicals, peptides, and recombinant proteins** | | |
| Dulbecco’s modified Eagle Medium | Gibco | Cat#12100046 |
| Opti-MEM Reduced Serum Medium | Gibco | Cat#31985070 |
| Minimum Essential Medium | Gibco | Cat#61100053 |
| Fetal Bovine Serum, heat inactivated | Gibco | Cat#16140071 |
| Penicillin-Streptomycin-Amphotericin B | MP Biomedicals | Cat#ICN1674049 |
| GlutaMAX™ | Gibco | Cat#35050-061 |
| Trypsin From Bovine Pancreas (TPCK Treated) | Sigma Aldrich | Cat#T1426 |
| Poly-L-lysine | Sigma Aldrich | Cat#P9155 |
| Polybrene | Merck | TR-1003-G |
| TRIzol™ Reagent | Thermo Fisher | Cat#15596018 |
| Phosphate Buffered Saline (10x) | MP Biomedicals | Cat#162528 |
| 4x Laemmli Sample Buffer | Bio-Rad | Cat#1610747 |
| Skimmed Milk | Sigma Aldrich | Cat#70166 |
| Tween 20 | Sigma Aldrich | Cat#P1379 |
| Xylazine | Indian Immunologicals Ltd. | Cat#21 |
| Ketamine | Bharat Parenterals Limited | N/A |
| 2-Picolinic acid | Sigma Aldrich | Cat#P42800 |
| Chloroquine diphosphate salt | Sigma Aldrich | Cat#C6628 |
| Ammonium chloride | Fisher Scientific | Cat#21405 |
| Octadecyl Rhodamine B Chloride (R18) | Invitrogen | Cat#O246 |
| Crystal Violet | MP Biomedicals | Cat#152511 |
| Formalin | Sigma Aldrich | Cat#F8775 |
| DAPI for nucleic acid staining | Sigma Aldrich | Cat#D9542 |
| ProLong™ Diamond Antifade Mountant | Invitrogen | P36970 |
| Lipofectamine 2000 transfection reagent | Invitrogen | Cat#11668019 |
| Wheat Germ Agglutinin, Alexa Fluor™ 488 Conjugate | Invitrogen | Cat#W11261 |
| Transferrin Alexa Fluor 647 Conjugate | Invitrogen | Cat#T23366 |
| DNase I (RNase-free) | New England BioLabs | Cat# M0303S |
| RNase A, DNase and protease-free | Thermo Scientific | Cat#EN0531 |
| SalI-HF | New England BioLabs | Cat#R3138S |
| Carbon Type-B, 300 mesh, Copper | Ted Pella | Cat#0813 |
| Uranyl Acetate | Ted Pella | Cat#19481 |
| TritonX100 | Sigma Aldrich | Cat#T8787 |
| Oxoid Agar | Oxoid Limited | Cat#LP0028 |
| DEAE-Dextran hydrochloride | Sigma Aldrich | Cat#D9885 |
| Sodium bicarbonate | MP Biomedicals | Cat#194553 |
| Sodium chloride | Sigma Aldrich | Cat#S3014 |
| HEPES (free acid) | Promega | Cat#H5303 |
| Magnesium chloride anhydrous | HiMedia | Cat#MB237 |
| Polyethylene glycol Mol Wt. 8000 | Sigma | Cat#P-2109 |
| Chloroform | Qualigens | Cat#Q12305 |
| Luria broth | HiMedia | Cat#M575 |
| Middlebrook 7H10 agar | Sigma Aldrich | Cat#M0303 |
| ADC growth supplement | HiMedia | Cat#FD019 |
| Tween 80 | Fisher Scientific | Cat#YBP338500 |
| Calcium Chloride | Fisher Scientific | Cat#Q12135 |
| Glycerol | Fisher Scientific | Cat#Q24505 |
| Agarose, low gelling temperature | Sigma-Aldrich | Cat#A9414 |
| **Critical commercial assays** | | |
| AgPath-ID™ One-Step RT-PCR kit | Applied Biosystems | Cat#AM1005 |
| alamarBlue™ Cell Viability Reagent | Invitrogen | Cat#DAL1025 |
| Clarity Western ECL Substrate | Bio-Rad | Cat#1705061 |
| Firefly luciferase assay kit | Promega | Cat#E4550 |
| Dual-Luciferase Reporter Assay System | Promega | Cat#E1980 |
| **Experimental models: Cell lines** | | |
| HEK293T-ACE2 | BEI Resources, NIAID, NIH | NR-52511 |
| Vero E6 | ATCC | CRL-1586 |
| HEK293T | NCCS, Pune, India | N/A |
| A549 | NCCS, Pune, India | N/A |
| MDCK | NCCS, Pune, India | N/A |
| HeLa | ATCC | CCL-2 |
| Calu-3 | ATCC | HTB-55 |
| **Experimental models: Organisms/strains** | | |
| Syrian Golden Hamster | Biogen laboratory animal facility | N/A |
| BALB/c mice | Central Animal Facility, Indian Institute of Science | N/A |
| **Oligonucleotides** | | |
| SARS-CoV-2 N1 Primers | Merck | VC00021N |
| SARS-CoV-2 N1 Probe | Merck | VC00023N |
| **Recombinant DNA** | | |
| Vector pHDM Containing the SARS-Related Coronavirus 2, Wuhan-Hu-1 Spike Glycoprotein, | BEI Resources, NIAID, NIH | NR-52514 |
| SARS-Related Coronavirus 2, Wuhan-Hu-1 Spike D614G-Pseudotyped Lentiviral Kit  Lentiviral Backbone, Luc2; ZsGreen | BEI Resources, NIAID, NIH | NR-52516 |
| SARS-Related Coronavirus 2, Wuhan-Hu-1 Spike D614G-Pseudotyped Lentiviral Kit  Helper plasmid, Gag; pol | BEI Resources, NIAID, NIH | NR-52517 |
| SARS-Related Coronavirus 2, Wuhan-Hu-1 Spike D614G-Pseudotyped Lentiviral Kit  Helper plasmid, Tat1b | BEI Resources, NIAID, NIH | NR-52518 |
| SARS-Related Coronavirus 2, Wuhan-Hu-1 Spike D614G-Pseudotyped Lentiviral Kit  Helper plasmid, Rev1b | BEI Resources, NIAID, NIH | NR-52519 |
| pAdDeltaF6 | Gift from James M. Wilson | Addgene plasmid # 112867, RRID:Addgene_112867 |
| pRepCap6 | Gift from David Russell | Addgene plasmid # 110770 ; RRID:Addgene_110770 |
| pAAV-CAG-GFP | Gift from Edward Boyden | Addgene plasmid # 37825 ; RRID:Addgene_37825 |
| pCB3/T7 | Kind gift from Prof. Frank van Kuppeveld | (van Ooij et al., 2006) |
| IAV Mini replicon Plasmids (PA, PB1, PB2, NP, NP-Luciferase) | Kind Gift from prof. Adolfo Garcia-Sastre (ISMMS, NY) | (Bortz et al., 2011) |
| pRLTK | Promega | E2231 |
| **Software and algorithms** | | |
| BioRender | BioRender | https://biorender.com/ |
| ImageJ/Fiji | (Schindelin et al., 2012) | https://fiji.sc/ |
| QuantStudio Design and Analysis Software v1.5.1 | Applied Biosystems | https://www.thermofisher.com/in/en/home/global/forms/life-science/quantstudio-3-5-software.html |
| CytExpert Acquisition and Analysis Software Version 2.3 | Beckman Coulter | https://www.mybeckman.in/flow-cytometry/instruments/cytoflex/software |
| GraphPad Prism 8.4.3 | GraphPad Software | https://www.graphpad.com/scientific-%20software/prism/ |
| Magellan | Software V7.1 SP1 | https://lifesciences.tecan.com/software-magellan |
| **Others** | | |
| Transmission Electron Microscope | Thermo Scientific | Talos L120C |
| CytoFLEX flow cytometer | Beckman Coulter | A00-1-1102 |
| Confocal multiphoton microscope | Zeiss | Zeiss LSM 880 |
| EVOS™ M5000 Imaging System | Invitrogen | AMF5000 |
| Ultracentrifuge | Beckman Coulter | L8-70M |
| ChemiDoc™ MP Imaging System | BIO RAD | 12003154 |
| Quantstudio 5 Real-Time PCR Instrument 384-well Block) | Applied Biosystems | A28135 |
| TECAN Infinite 200-PRO multiplex reader. | TECAN | https://lifesciences.tecan.com/plate_readers/infinite_200_pro |
| 7205 UV/Visible scanning spectrophotometer | Jenway | http://www.jenway.com/product.asp?dsl=9171 |
| Tecan Spark multi-mode plate reader | Tecan | https://lifesciences.tecan.com/multimode-plate-reader |
| Formvar/carbon-covered 300 mesh copper grid | Ted Pella | Cat#01753-F |
| PVDF membrane | Immobilon-P; Merck | Cat#IPVH00010 |
| Amicon Ultra-0.5 Centrifugal Filter Unit | Merck-Millipore | Cat#UFC510008 |
| Whatman® UNIFLO® 25 syringe filters, pore size 0.45 μm | Sigma Aldrich | Cat#9913-2504 |
| Whatman® qualitative filter paper, Grade 1 | Sigma Aldrich | Cat#WHA1001125 |
